# Generative machine learning unlocks the first proteome-wide image of human cells

**DOI:** 10.64898/2026.03.31.715748

**Authors:** Huangqingbo Sun, Konstantin Kahnert, Jan N. Hansen, William Leineweber, Mingyang Li, Wanyue Feng, Frederic Ballllosera Navarro, Ulrika Axelsson, Wei Ouyang, Emma Lundberg

## Abstract

The spatial organization of proteins within cells governs virtually all cellular functions, yet current imaging can simultaneously visualize only tens of proteins, orders of magnitude below the thousands populating a single human cell. Here we present *ProtiCelli*, a deep generative model that simulates microscopy images for 12,800 human proteins from just three cellular landmark stains. Trained on 1.23 million Human Protein Atlas images, *ProtiCelli* outperforms existing methods in reconstruction accuracy and textural fidelity, and generalizes to unseen cell types and drug perturbations. Simulated images preserve hierarchical subcellular organization, recapitulate known protein–protein interaction landscapes, and resolve compartment-specific functions of moonlighting proteins at single-cell resolution. Remarkably, the model infers drug-induced changes in protein expression and localization from cell morphology alone, predicts cell cycle stage without dedicated markers, and enables unsupervised segmentation of subcellular compartments and spatial decomposition of gene sets into functional regions. We leverage *ProtiCelli* to generate *Proteome2Cell*, a dataset of 30.7 million simulated images spanning 2,400 ”virtual cells” across 12 human cell lines, enabling hierarchical single-cell models that distinguish conserved from dynamic protein architectures. Integrated into the Human Protein Atlas, *Proteome2Cell* democratizes exploration of these virtual cells. By computationally bridging the experimental scalability gap, *ProtiCelli* establishes a foundation for spatial virtual cell modeling.

## Main

Cellular structure and function are driven by the expression and localization of proteins, many of which dynamically relocalize during signal transduction, trafficking, or structural reorganization [1–3]. The disruption of protein localization underpins numerous diseases such as neurodegeneration, cancer, and metabolic disorders [4]. Remarkably, an estimated one-sixth of pathogenic coding variants lead to protein mislocalization [5]. The spatial distribution of the intracellular proteome within and across cell populations is therefore essential for a nuanced understanding of cellular behavior and disease states.

Large-scale initiatives such as the Human Protein Atlas (HPA) have systematically mapped protein expression and subcellular localization across human cell types using immunofluorescence (IF) microscopy, imaging one protein at a time alongside reference markers that delineate cellular landmarks [6, 7]. These efforts reveal that more than half of all human proteins localize to multiple subcellular compartments, often performing different functions in the different compartments. This adds a layer of spatial complexity beyond gene and protein expression levels. However, current experimental approaches face fundamental scalability constraints [8]. Although recent advances in multiplexed spatial proteomics enable simultaneous visualization of multiple proteins within individual cells [9], even state-of-the-art methods are limited to an average of 37 proteins [10]. This represents several orders of magnitude below the thousands of proteins in a human cell [6, 11]. Additionally, substantial variation in protein expression and localization between individual cells [1, 7, 12, 13] means that a massive amount of experimentation would be needed to image every protein in every cell type.

Generative modeling offers a computational solution to the challenge. Two main categories exist. The first category, bottom-up modeling, comprises two strands. Datadriven statistical methods learn parametric generative models of cell and organelle geometry directly from images, capturing cell-to-cell variation in shape, number, and spatial arrangement, as in CellOrganizer [14–18]; physics-based methods simulate subcellular organization from mechanistic first principles [19–22]. Both are compact and interpretable but have limited capacity to model complex spatial patterns. The second category is end-to-end modeling of protein images using deep learning approaches without explicit parameterization, which offers greater scalability and application flexibility [23–27]. Such state-of-the-art models include (1) the *Statistical Cell* model, a variational autoencoder that predicts 24 subcellular structures [25]; (2) the *PUPS* model, a U-Net-based image-inpainting model that uses protein language model embeddings as conditioning to predict the subcellular localization of proteins [26]; and (3) the *CELL-Diff* model, a unified diffusion model enabling bidirectional transformations between protein sequences and microscopy images [27]. *PUPS* and *CELL-Diff* are both trained on the HPA image collection and provide near proteome-wide prediction of subcellular protein localization.

Despite recent progress, fundamental limitations persist in deep learning models designed to virtually stain cells. Current models are systematically biased toward nuclear and cytosolic proteins because they are conditioned on limited cellular landmarks, thereby under-representing other compartments. Moreover, the field lacks comprehensive evaluation frameworks capable of assessing model performance from multiple complementary biological perspectives. To date, model development has emphasized technical benchmark evaluation. The few biological questions that have been addressed with prior virtual staining models related cell morphology to organelle positioning [25] and supported *in silico* screening and hypothesis generation [26, 27]. The systematic use of virtual staining to generate biological insight at scale remains an emerging effort.

Here we present *ProtiCelli*, a deep generative model that simulates immunofluorescence (IF) protein staining images based on cellular landmarks, trained on images from the HPA (Figure 1). *ProtiCelli* outperforms existing models and generalizes to unseen cell lines and perturbed cell states. Additionally, we establish a framework for evaluating virtual staining models through diverse biology-focused applications, such as predicting protein co-localization, cellular states (i.e., the cell cycle), drug-induced protein localization/intensity changes, or subcellular compartment segmentation. We show *ProtiCelli* ’s capability to enhance image databases like the HPA with virtual stains for experimentally unmeasured protein-cell-line combinations. Lastly, we simulate proteome-scale images of single cells covering the majority of intracellular proteins (*n*=12,800), a multiplexing depth exceeding current experimental methods by two orders of magnitude. Such simulated images enable the study of single-cell architecture in unprecedented complexity. These capabilities establish *ProtiCelli* as a foundational step towards building comprehensive virtual cell models capable of simulating cellular behavior *in silico* [28, 29], opening new avenues in functional genomics and precision medicine.

**Fig. 1:**
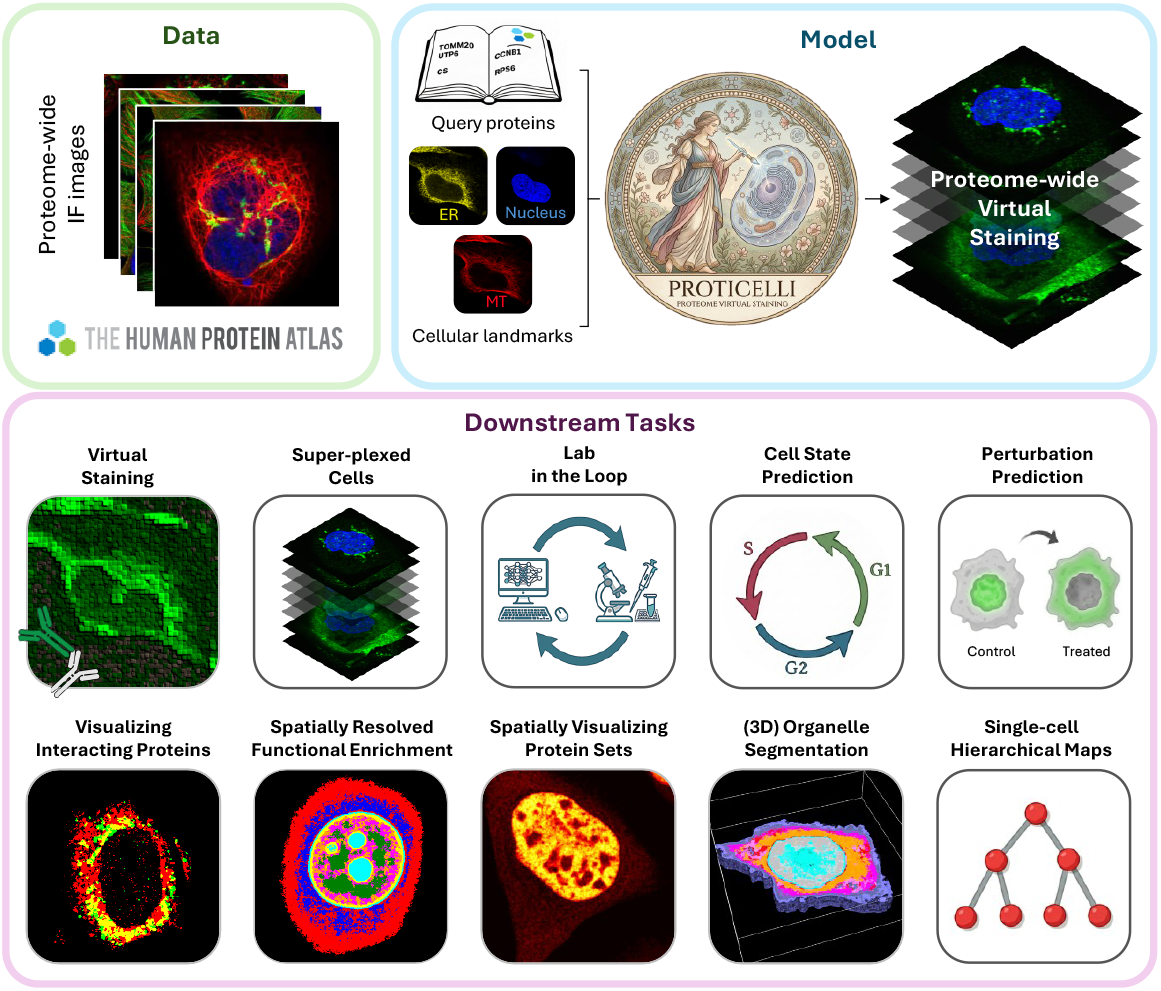
***ProtiCelli* study overview.** *ProtiCelli* was trained on near-proteome-wide single-cell-cropped immunofluorescence images from the Human Protein Atlas to simulate high-resolution fluorescent microscope images of 12,800 proteins conditioned on cellular landmarks (nucleus, endoplasmic reticulum (ER), and microtubules (MT)), which enables several downstream applications.

### ProtiCelli design

*ProtiCelli* simulates IF protein staining on single-cell image crops that provide three cellular landmark channels: nucleus, endoplasmic reticulum (ER), and microtubules (Figure 1). Because these landmark channels inform but do not fully specify protein localization, we formulated virtual staining as conditional generative modeling, learning the distribution of plausible protein staining patterns rather than a single deterministic prediction.

We implemented *ProtiCelli* as a conditional latent diffusion model [30–32] (Figure S1) using the pretrained variational autoencoder from Stable Diffusion 3.5 [33] and a Diffusion Transformer (DiT-L) architecture [34] (Methods). We selected a diffusion model because it learns the full conditional distribution, produces sharp and diverse samples, and trains stably. We chose the transformer-based approach over convolutional U-Nets to enable self-attention, thereby capturing long-range spatial dependencies and scaling effectively to proteome-scale data [35, 36]. The model was trained using the Elucidating Diffusion Models framework for time-efficient image generation [37] to broaden applicability to whole-proteome simulation.

We trained *ProtiCelli* on the near proteome-wide HPA image collection, from which we created approximately 1.23 million single-cell cropped images representing 12,800 human proteins across 39 cell lines (Table S1). Each image has been expert-annotated for protein localization in any combination of 40 organelles and subcellular structures. The cell lines exhibit diverse cellular morphologies, are derived from over 20 tissues, and have both cancerous and non-cancerous origins, including epithelial, mesenchymal, myeloid, and endothelial origins (Figure S2A). Images were split into training (*HPA-training*, 1.15 million images) and testing (*HPA-testing*, 80,000 images) sets, stratified by cell line and organelle (Figures S2B-E) (Methods).

We made two key design choices while constructing *ProtiCelli* that set it apart from other simulated IF protein staining models. First, *ProtiCelli* is conditioned with learnable per-protein embeddings rather than protein-sequence embeddings from language models. Although sequence-based conditioning enables generalization to unseen proteins [26, 27], we observed that it substantially reduced image quality (FID increase from 6.46 to 38.27), in line with recent work demonstrating that current protein language models fail to capture localization signals [38]. Our fixed-protein-vocabulary approach prioritizes accurate spatial representation of proteins while still covering 12,800 proteins, the majority of all intracellular human proteins. Second, to enable generalization to unseen cell lines, we stochastically withheld cell-line conditioning during training (Methods). The *ProtiCelli* model generates high-resolution (512×512 pixels) single-cell-cropped images for all 12,800 trained proteins based on providing the three cellular landmark channels (Video S1, Figure 2A).

**Fig. 2:**
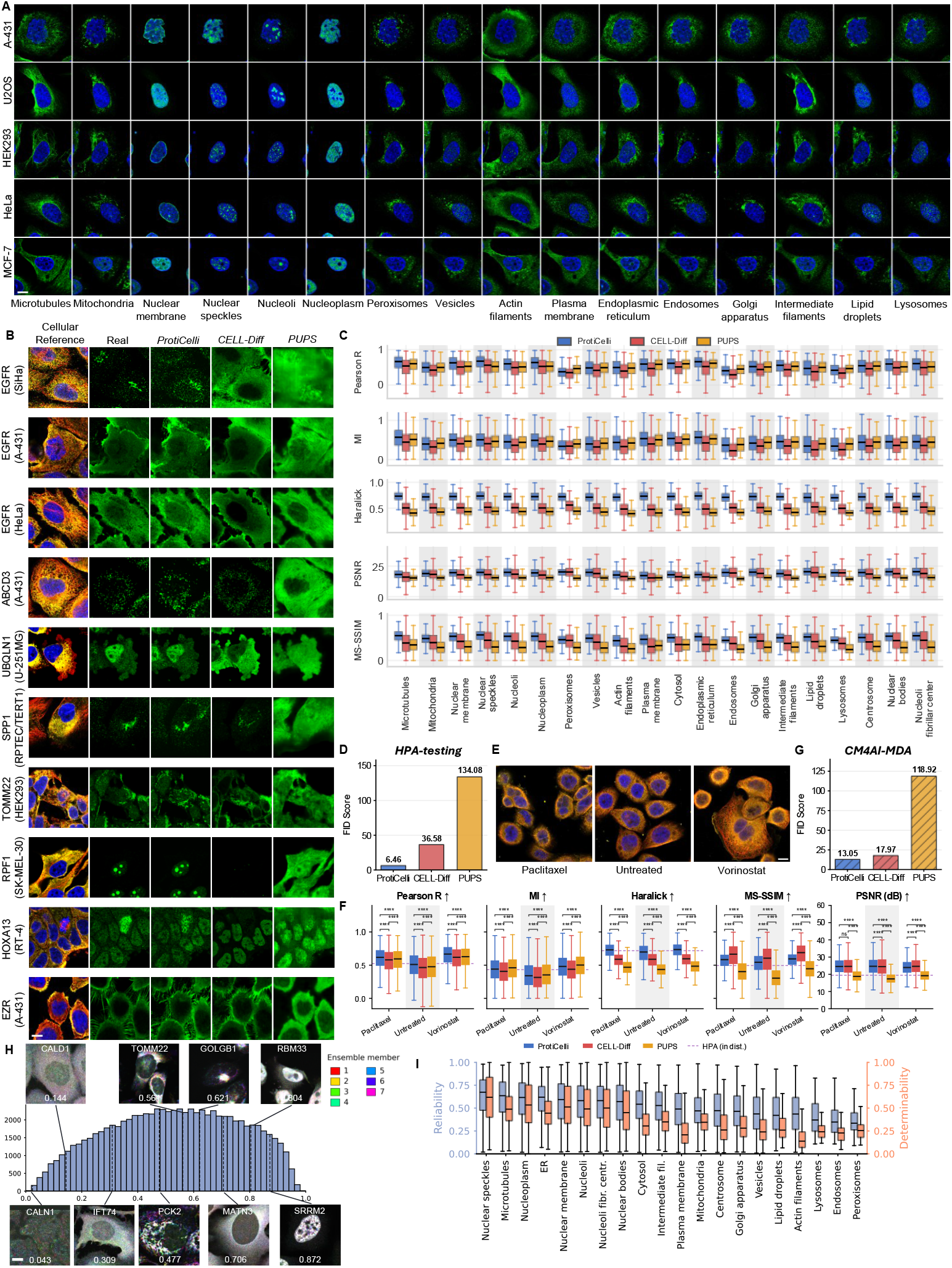
Benchmarking *ProtiCelli* (continued). (**A**) *ProtiCelli* -simulated immunofluorescence images of major organelles and cellular structures. (**B**) Representative images simulated by *ProtiCelli*, *CELL-Diff* and *PUPS*. In SiHa cells, EGFR exhibits localization to the Golgi apparatus rather than its typical plasma-membrane localization. This cell line-specific localization is captured only by *ProtiCelli*. Cellular- reference images used as model inputs are the HPA landmark channels: nucleus (DAPI, blue), endoplasmic reticulum (yellow) and microtubules (red). (**C**, **D**) Performance of *ProtiCelli*, *PUPS* and *CELL-Diff* on held-out test images, evaluated using metrics of pixel-level accuracy (Pearson’s *r*, mutual information (MI), peak signal-to-noise ratio (PSNR)), structural similarity (multiscale structural similarity index measure (MS-SSIM)), textural similarity (Haralick-feature similarity), and semantic image quality (Fŕechet Inception Distance (FID); lower values indicate better performance; averaged across all localization classes). (**E**) Representative immunofluorescence images from the *CM4AI-MDA* dataset, comprising MDA-MB-468 cells under control and drug-treatment conditions. Only the landmark channels are shown, colored as in **B**. (**F**, **G**) Performance on the *CM4AI-MDA* dataset, evaluated using the metrics in **C**, **D**. *ProtiCelli* achieves the best overall performance (arrows), comparable to its in-distribution performance (dashed line in **F**: mean *ProtiCelli* score in **C**). (**H**) Histogram of reliability scores across the *HPA-testing* set. Seven of the 20 ensemble members, selected at random, are pseudo-colored and overlaid, such that spatially concordant predictions sum to white and divergent predictions appear as separate hues. This illustrates how the reliability score (shown per panel) tracks cross-sample agreement: high-reliability proteins (e.g. SRRM2, 0.87) render as clear white structure, low-reliability proteins (e.g. CALN1, 0.02) fragment into scattered color. (**I**) Reliability (left axis, blue) and determinability (right axis, orange) by organelle in *HPA-testing*, ordered by mean reliability. Boxes indicate the median and interquartile range; whiskers extend to the data range excluding outliers. All scale bars, 10 µm.

### Subcellular patterns and cell topology

Human cells are hierarchically and spatially organized, from membrane organelles such as mitochondria to biomolecular condensates such as nucleoli [1, 39]. Useful generative cell models must simulate fine structures (e.g., actin filaments, mitochondria, nuclear speckles), diffuse structures (nucleoplasm or cytosol localization), and multi-localizing proteins [1]. By eye, *ProtiCelli* -simulated images appeared realistic across diverse subcellular patterns and cell lines (Figure 2A). To quantitatively evaluate *ProtiCelli* ’s performance, we used the reference channels (nucleus, ER, microtubules) from each cell in *HPA-testing* to simulate the respective protein channel and compared it to the corresponding ground-truth protein channel. We compared results against those obtained with the state-of-the-art models *CELL-Diff* [27] and *PUPS* [26]. *ProtiCelli* outperformed *CELL-Diff* and *PUPS* visually, generating images of realistic and more accurate subcellular structures (Example: Figures 2B and S3), and captured cell-line-specific localization patterns that other models missed. For example, EGFR localizes to the Golgi apparatus in SiHa cells but to the plasma membrane in most other cell lines (U2OS, A-431). Whereas *CELL-Diff* and *PUPS*uniformly predicted plasmamembrane localization, *ProtiCelli* correctly simulated Golgi localization in SiHa cells (Figure 2B). This highlighted a consistent strength of *ProtiCelli* for multi-localizing proteins (Figure S4A).

Quantitative metrics across reconstruction accuracy, textural fidelity, and perceptual quality (Figure 2C) confirmed the visual observations that *ProtiCelli* outperformed *CELL-Diff* and *PUPS*. For pixel-level reconstruction, *ProtiCelli* achieved superior or matching Pearson correlations and mutual information scores for all organelles except the vesicular compartments: peroxisomes, endosomes, and lysosomes. All models performed poorly on vesicles, likely due to the weak spatial correlation of vesicle positioning with the three cellular landmarks. In contrast, pixel-level performance was particularly strong for nuclear structures (e.g., nuclear membrane, nuclear speckles, nucleoli), suggesting that the texture of nuclear staining is highly informative for these patterns. *ProtiCelli* excelled at recapitulating textural features (Haralick, MS-SSIM, Figure 2C). Overall fidelity and cell topology representation was also substantially superior for *ProtiCelli* (Figure 2D). The UNet-based *PUPS* architecture, trained with a pixel-wise MSE reconstruction objective and an auxiliary subcellular localization classification loss, is known to produce spatially over-smoothed outputs [40, 41], explaining lower textural performance. The performance gap with *CELL-Diff* likely reflects the per-protein-vocabulary design, higher output resolution of *ProtiCelli*, and the greater generative capacity of the EDM framework combined with the DiT-L backbone.

In general, *ProtiCelli* demonstrated superior spatial accuracy compared to stateof-the-art models, correctly positioning proteins within their appropriate cellular compartments with high textural fidelity. Applying multiple quantitative metrics that assess different aspects of image quality, we advanced beyond standard benchmarking of virtual stains, especially for fine tubular, sheet-like, and vesicular structures. Our tests confirm that *ProtiCelli* effectively learned to represent spatial patterns of diverse organelles and subcellular structures without explicit supervision of subcellular protein localization during training.

### Unseen cell line and cell states

Staining every protein in every cell type and every perturbation condition is infeasible. Virtual staining models promise to perform *in silico* experiments to prioritize real experiments effectively, requiring the models to generalize to unseen contexts. To evaluate how *ProtiCelli* generalizes to new biological contexts, we explored performance on an independently acquired dataset unseen by any of the models: *CM4AI-MDA* [42]. The *CM4AI-MDA* dataset consists of 112,410 cells from the MDA-MB-468 triplenegative breast cancer cell line, treated with the anti-cancer drugs Paclitaxel or Vorinostat, or vehicle control (untreated) (Figure 2E), and comprises IF images covering 453 proteins. Cell line or perturbation information was not provided to *ProtiCelli*, so a drug treatment can shape prediction only through changes in the landmark channels. *ProtiCelli* maintained strong performance on MDA-MB-468 cell images across all metrics, and retained its advantage over *PUPS* and *CELL-Diff* (Figures 2F and G), in particular when focusing on texture (e.g., Haralick, Figure 2F). The robust generalization likely reflects the morphological diversity of the training data, providing a breadth of cell shapes in cancer and non-cancer cell lines. This test shows that *ProtiCelli* generalizes to both unseen cell lines and experimentally perturbed conditions.

### Reliability-aware staining simulations

*ProtiCelli* can simulate the expression patterns of 12,800 proteins. However, the accurate positioning of a staining pattern in a cell may depend on the breadth of cellular contexts in which *ProtiCelli* has seen each specific protein and on how informative the reference markers are for positioning a specific subcellular pattern. For example, we observed that *ProtiCelli* predicted microtubule staining (which is one of the reference channels) with much higher pixel-based accuracy than vesicle staining (Figure 2C).

To estimate the reliability of simulated staining patterns we introduced an ensemble-agreement approach (Methods): for each cell and protein, we generate multiple (*N* = 20) independent predictions of the protein channel and measure agreement across the ensemble. Applying this approach to the *HPA-testing* data, we observed a wide distribution of reliability scores (Figure 2H) and a systematic dependence on subcellular compartments (Figure 2I): nucleus-associated structures and microtubules scored highest, whereas vesicular structures scored lowest, consistent with their more stochastic localization and a weaker correlation with reference channels. Reliability strongly correlated with an independent measure of how well each compartment is determined from the reference markers in ground-truth images (Spearman’s *ρ* = 0.827; Figure 2I, S4B, C; Methods). Correlating the reliability scores with established image-similarity metrics across *HPA-testing* revealed a strong correlation with pixel-level fidelity in particular (*r* = 0.84 for Pearson *r*; MI: *r* = 0.56, MS-SSIM: *r* = 0.47, Haralick: *r* = 0.24, PSNR: *r* = −0.10; Methods).

These results show that the reliability score reflects biologically intrinsic challenges for predicting subcellular compartment localization from a limited set of reference markers, providing users with a per-protein indicator of positional credibility for each of the 12,800 simulated protein staining patterns. Importantly, a low reliability score does not indicate an unrealistic staining pattern, but rather that the exact positioning of a plausible pattern is less certain. A simulated vesicle pattern, for instance, reliably predicts vesicular localization in a plausible region but not the exact vesicle coordinates, whereas microtubule or nuclear patterns are reliable down to the pixel.

### Proteome-wide single-cell images

After benchmarking image quality and reliability, we explored how simulated images could inform biological tasks. Because even state-of-the-art highly multiplexed staining approaches are limited to revealing a few to hundreds of proteins within individual cells, we explored applying *ProtiCelli* on a systems-biology level, virtually staining 200 cells from 12 cell lines for all 12,800 proteins that *ProtiCelli* can simulate. This dataset, called *Proteome2Cell*, comprises 30,720,000 simulated images covering osteosarcoma (U2OS), squamous cell carcinoma (A-431), glioblastoma (U-251MG), breast adenocarcinoma (MCF-7), cervical adenocarcinoma (HeLa), hepatocellular carcinoma (Hep-G2), colorectal adenocarcinoma (CACO-2), prostate adenocarcinoma (PC-3), rhabdomyosarcoma (Rh30), neuroblastoma (SH-SY5Y), transformed embryonic kidney (HEK293), and normal skin fibroblast (BJ) cells. *Proteome2Cell* provides the first comprehensive view of the majority of intracellular proteins within individual cells, enabling systematic exploration of proteome-wide spatial organization (Video S1).

### Hierarchical protein organization

We evaluated *Proteome2Cell* by encoding the simulated images with the *SubCell* foundation model [43], which captures single-cell patterns of subcellular protein localization and cell morphology, and comparing them to embeddings from ground-truth (real) HPA images. To control for morphology, sampling depth, and domain shifts, we analyzed cell lines separately, matching cell counts before averaging single-cell embeddings into one vector per protein and centering the final representations (Methods).

When visualized using a joint UMAP projection, real and simulated representations overlapped and organized similarly by localization (Figure 3A, Figure S5A). Quantitatively, within the high-dimensional space itself, 59-66% of a protein’s 15 nearest neighbors shared its source (real or simulated; 50% expected if fully intermixed). Embeddings of simulated images also achieved 84% of the neighborhood localization purity of real images. We conclude the simulated images mostly resemble the real atlas. To confirm the simulated images carry biological information beyond shared occupancy, we predicted each protein’s location using its five nearest neighbors in the embeddings of real images (Figure 3B). Because SubCell encodes both localization and morphology, we established a morphology-only baseline by blanking the protein channel. This reduced performance to chance (macro F1 of 0.04), confirming morphology alone provides negligible predictive signal. Evaluated against the real-to-real upper bound, simulated images retained 90% of the top-1 accuracy (45.2% vs. 50.5%) and 69% of the macro F1 score (0.17 vs. 0.25) across all 12 cell lines. This indicates that *ProtiCelli* -simulated images largely reflect the subcellular localization pattern of the individual protein they depict.

**Fig. 3:**
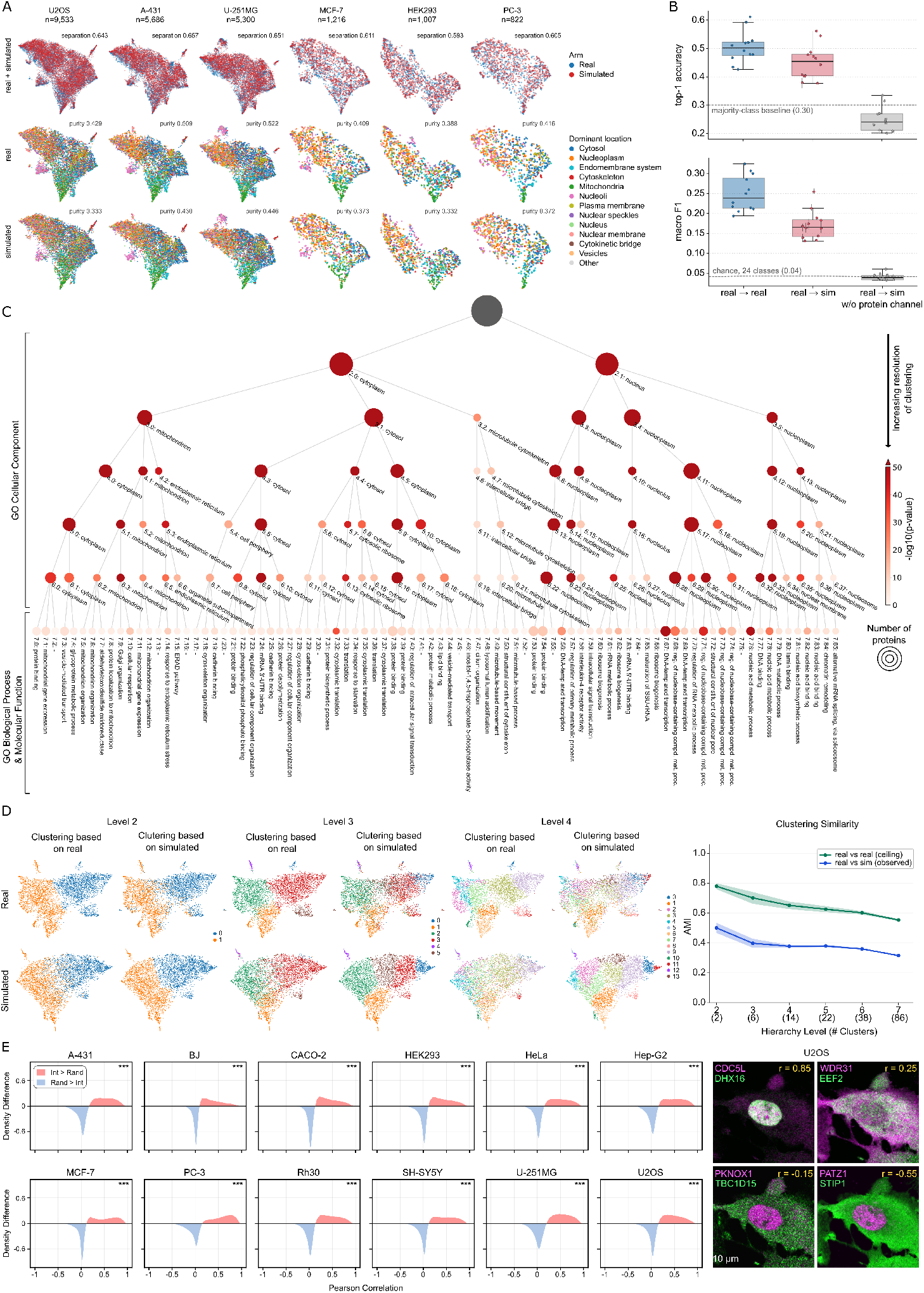
Learned representations (continued). (**A**) UMAP projections of *SubCell* embeddings for the six HPA cell lines with the most stained proteins; the remaining six are shown in Fig. S5A. Each point represents one protein. Embeddings of real and simulated images were mean-centered and jointly projected for each cell line (Methods). Columns denote cell lines; the top row shows embeddings colored by image source, and the lower rows show embeddings of real and simulated images separately in the same projection, colored by primary subcellular location. Separation and purity were calculated using the full 1,536-dimensional embeddings. (**B**) Accuracy of subcellular-location prediction from protein embeddings using the majority location among the five most similar proteins in a reference set of real images. Inputs comprised images containing real or *ProtiCelli* -simulated protein channels, or images with the protein channel blanked (morphology only). *Top*, classification accuracy; *bottom*, macro-averaged F1 score. Each point represents one cell line (*n* = 12) and the mean of three simulated-cell sampling repeats. Whiskers for the real–simulated condition denote the range across repeats (Methods); boxes denote the median and interquartile range. Dashed lines indicate majority-class (0.30) and chance-level (0.041 for 24 classes) performance. (**C**) Hierarchical map of subcellular organization constructed from embeddings of simulated images using Leiden clustering at progressively finer resolutions, annotated with enriched GO terms. Node size reflects cluster size; color intensity reflects enrichment significance. (**D**) Agreement between real and simulated maps in U2OS cells. *Left*, UMAP projections colored by clusters at hierarchy levels 2–4, with clustering performed in the high-dimensional embedding space. Rows denote embeddings of real and simulated images; columns denote hierarchy level and the dataset used to define cluster colors. *Right*, adjusted mutual information (AMI) between clusterings across hierarchy levels. Curves compare real and simulated images (blue) or two independent sets of real images (green). Data points and bands are mean ±1 s.d. over 30 runs. (**E**) *Left*, differences between the distributions of pixel-level Pearson correlations for interacting and localization-matched non-interacting protein pairs across 12 cell lines. Positive values (red) indicate higher correlations among interacting pairs, whereas negative values (blue) indicate higher correlations among localization-matched non-interacting pairs.Significance was assessed using a two-sided Mann–Whitney *U* -test; *** *p* ¡ 0.005. *Right*, representative *ProtiCelli* simulated images of known interacting and unrelated protein pairs in U2OS cells, shown in magenta and green. Scale bar, 10 µm.

### Fidelity across cellular hierarchies

A defining feature of cellular architecture is hierarchical organization: major compartments (cytoplasm, nucleus) subdivide into organelles (mitochondria, Golgi, ER) or phase-separated regions (nucleoli), which further subdivide into protein assemblies. To evaluate whether *ProtiCelli* preserves this organization, we constructed a multi-scale cell map by multi-resolution Leiden clustering of the *SubCell* -embeddings of *Proteome2Cell* ’s U2OS cells and compared it to a map from real images [43] (Figure 3C and Figure S5B). The hierarchy derived from simulated images mirrored that from real cells: primary nuclear-cytoplasmic separation followed by resolution of the nucleolar, nucleoplasmic, and nuclear membrane subdomains, and by separation of the cytoplasmic compartments like mitochondria or the cytoskeleton. Simulated and real maps agreed most at the nucleus–cytoplasm split (AMI = 0.50) and less with depth (0.32 across 86 clusters) (Figure 3D); the same decline appeared when comparing two independent sets of real images (0.78 to 0.55). Crucially, the simulated map captured a consistent fraction of the real-to-real baseline across all resolutions (57–64%), demonstrating that *ProtiCelli* follows the empirical limits of real data rather than degrading at finer scales. This indicates that *ProtiCelli* -simulated images reproduce proteome organization from whole compartments down to functionally coherent protein groups. In contrast to the HPA, which pictures one protein per cell, *Proteome2Cell* simulates all proteins in the same cells. This enables us to study how proteins are arranged relative to one another. To validate this, we tested whether known physically interacting proteins are placed together in *Proteome2Cell* images by measuring pixel-wise spatial co-localization, as established for real images [43, 44]. Across 309,411 high-confidence STRING interactions (score *>*850), interacting pairs showed significantly higher spatial correlation than a localization-matched null (same primary compartments; whole-cell region) in all 12 cell lines (Figure 3E; Methods). This indicates that *ProtiCelli* not only recapitulates compartment localization but also spatial arrangements that reflect physical interactions of proteins.

### Spatially-resolved protein interaction

A requirement for physical interaction of two proteins is that they are co-localizing. More than half of all human proteins localize to multiple subcellular compartments [7], and many of these proteins perform different functions in different compartments, a phenomenon known as protein ”moonlighting” [1]. *ProtiCelli* simulations can help disentangle moonlighting protein function by resolving compartment-specific co-localization. As a proof of concept, we explored the known moonlighting enzyme P4HA2 [45]. In U-251MG cells, P4HA2 localizes to the ER and vesicles in some cells, and to the nucleoplasm in other cells (Figure 4A (red outlined images)) [45]. We simulated staining images for known interaction partners of P4HA2 to predict compartment-specific interaction landscapes. ER-localized collagen pathway partners (COL12A1, COL14A1, COL1A1, P4HB, COL18A1) co-localized with P4HA2 at the ER, while members of the CPSF complex involved in mRNA polyadenylation (FIP1L1, CPSF2, CPSF4, SYMPK, WDR33) co-localized with P4HA2 in the nucleoplasm. The compartment-specific segregation of P4HA2’s known interaction partners suggests distinct interaction contexts and nominates a nuclear moonlighting role in mRNA processing (Figure 4A). While spatial co-localization is only a prerequisite for, but not proof of, compartment-specific interaction, the P4HA2 example demonstrates *ProtiCelli* ’s capacity to resolve context-dependent localization patterns and generate testable hypotheses about compartment-specific protein function.

**Fig. 4:**
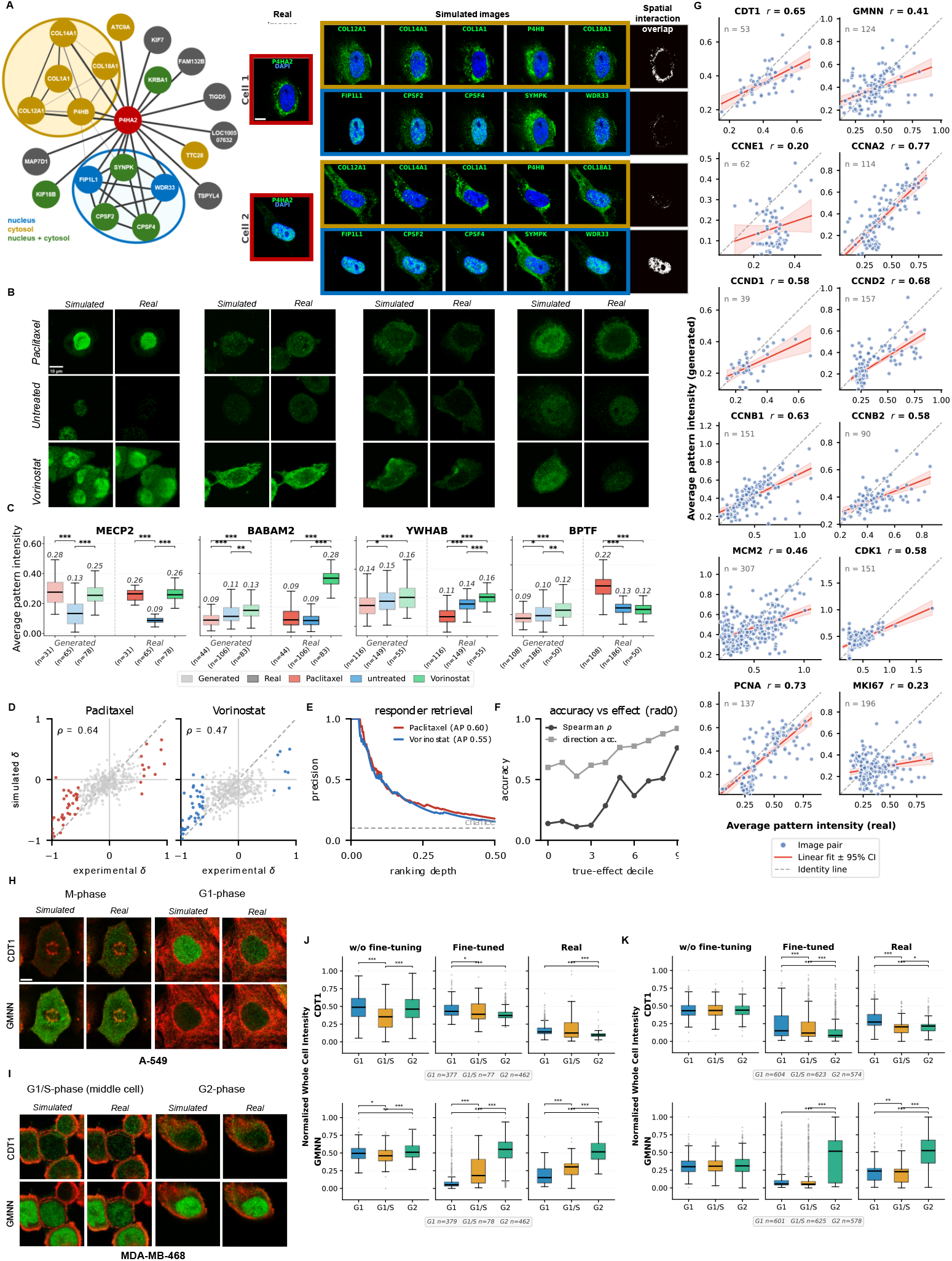
***ProtiCelli* captures protein interaction context, drug-induced expression variance, and cell-cycle-dependent dynamics.** (**A**) *Left*, protein- protein interaction network centered on P4HA2, showing interactions with proteins localized to the endoplasmic reticulum (ER) and nucleoplasm. *Right*, real P4HA2 images (red border) and *ProtiCelli* -simulated images of ten interaction partners in two representative cells, grouped by interaction context (gold, ER/collagen; blue, mRNA processing). The rightmost column shows binarized spatial overlap between P4HA2 and the mean projection of each partner group. (**B**) Representative real and *ProtiCelli* - simulated fluorescence microscopy images of MECP2, BABAM2, YWHAB and BPTF in Paclitaxel, untreated, and Vorinostat-treated cells. (**C**) Normalized intensities of the four proteins in **B** across treatment conditions in real and *ProtiCelli* -simulated images. Boxes show the median and interquartile range; annotated values indicate medians. Brackets indicate comparisons assessed using a Mann–Whitney *U* -test. **(D)** Experimental versus simulated Cliff’s *δ* for each of the 438 proteins measured under both Paclitaxel and Vorinostat treatment. The dashed line indicates *y* = *x*; colored points denote the 10% of proteins with the largest experimental effect magnitudes. Spearman correlation coefficients are indicated. **(E)** Precision for retrieving strong responders ranked by simulated effect magnitude, plotted against ranking depth over the top half of the ranking for each drug. The dashed line indicates the base-rate expectation (0.10); filled circles indicate a ranking depth of 10%. Average precision is shown in the legend. **(F)** Spearman correlation (*ρ* = 0.14 to 0.76) and directional agreement (0.60 to 0.92) between experimental and simulated effects within deciles of experimental effect magnitude. (**G**) Mean pattern intensity in real and *ProtiCelli* - simulated images for 12 cell-cycle-regulated proteins. Each point represents one cell; red lines and shading indicate linear fits and 95% confidence intervals, respectively, and dashed lines indicate *y* = *x*. Pearson’s *r* is reported for each protein. (**H**, **I**) Representative real and *ProtiCelli* -simulated immunofluorescence images of the FUCCI markers CDT1 and geminin (GMNN) in A-549 (**H**) and MDA-MB-468 (**I**) cells. Protein signal is shown in green; the microtubule reference from the corresponding real image, identical between real and simulated panels, is shown in red. (**J**, **K**) Simulated and measured intensities of CDT1 and geminin across cell-cycle stages (G1, G1/S and G2) before and after fine-tuning in A-549 (**J**) and MDA-MB-468 (**K**) cells. Significance was assessed using a Mann–Whitney *U* -test. All scale bars, 10 µm.

### Drug-response prediction

Genetic and small molecule perturbations often induce changes in protein expression, interaction, and localization [1, 5, 46]. Such changes, particularly protein mislocalization, can rewire signaling to drive disease or mediate therapeutic effects through loss or gain of interactions and function. Yet, systematic analysis of drug-induced protein responses remains experimentally challenging at the proteome scale and the current single-cell analyses are limited to cell morphology readouts [47–49]. Since *ProtiCelli* has learned the relationship between cell morphology and protein expression, we explored *ProtiCelli* ’s applicability for predicting drug-induced changes in protein abundance and localization from cellular morphology, without receiving the drug identity.

On the *CM4AI-MDA* dataset, *ProtiCelli* accurately predicted many expression and localization changes, such as increased nucleoplasmic signal of MECP2 under Paclitaxel and increased cytosolic signal of BABAM2 under Vorinostat treatment (Figures 4B, C, and Table S2). A failure example is BPTF, redistributing to the plasma membrane upon Paclitaxel treatment, which is not simulated by *ProtiCelli* (Figure 4B). Of note, BPTF is annotated exclusively to the nucleoplasm in the HPA and therefore its redistribution to the plasma membrane is unpredictable because this localization lies outside the training distribution (Figure S6A). *ProtiCelli* reproduced drug-induced expression changes (mean protein staining intensity) across proteins with above-chance accuracy (Table S2). Beyond intensity, simulated images also captured changes in protein spatial organization: radial redistribution, quantified by Cliff’s *δ*, correlated between experimental and simulated images (Spearman *ρ* = 0.64 for Paclitaxel and 0.47 for Vorinostat; Figure 4D; Table S3). Ranked by simulated response, proteins with the largest experimental responses were strongly enriched in the top group (average precision 0.60 for Paclitaxel, 0.55 for Vorinostat; chance 0.10; Figure 4E). Agreement of simulated and experimental effect increased with experimental effect magnitude (Spearman *ρ* = 0.76 and 92% directional agreement for top decile; Figure 4F), indicating that the strongest drug-induced changes were most reliably recovered by *ProtiCelli*.

Together, these results show that *ProtiCelli* recovers both responsive proteins and quantitative changes in their subcellular distribution during drug perturbation, despite never receiving the drug identity. The cellular morphology encodes rich molecular information about proteome spatial organization beyond what standard image profiling captures. The model likely learns these relationships from diverse HPA training data spanning cancer and primary cell types with varied morphologies (Figure S2A), enabling generative models to serve as tools linking morphological features to molecular states. However, *ProtiCelli* is limited to protein behaviors observed during training and cannot predict novel relocalization events absent from the HPA dataset. Training on expanded datasets, including perturbed cells, could improve the accuracy of drug response prediction and broaden the coverage of perturbation-induced protein behaviors.

### Cell-state prediction

Protein expression, localization, and function depend on the state of the cell, and the cell cycle represents a fundamental axis of dynamic cell states [13]. We investigated whether *ProtiCelli* could capture cell cycle-dependent expression dynamics by generating images of established cell cycle markers across all HPA cell lines. *ProtiCelli* correctly simulated relative expression patterns of many, but not all, tested cell cycle proteins, achieving a Pearson correlation coefficient of 0.587 between predicted and experimental intensities (*p <* 0.001, Figure 4G; additional examples in Figure S6B).

We further examined *ProtiCelli* simulations for the gold-standard FUCCI cell cycle markers CDT1 and GMNN [50]. Accurate simulation of these markers would enable computational cell cycle staging from reference morphology alone. We generated *FUCCI-CellCycle*, an HPA-style dataset of 574 images of cells stained with CDT1 and GMNN along with the three cellular reference channels (microtubules, ER, and DAPI) in A-549 (in-distribution; 322 images) and MDA-MB-468 (out-of-distribution; 252 images) cell lines. We simulated CDT1 and GMNN stains for these images with *ProtiCelli*, as exemplified in Figures 4H-I, and assigned each cell to its cell cycle stage using the ground-truth FUCCI marker staining. Under normal FUCCI dynamics, CDT1 declines and GMNN rises from G1 through G1/S to G2. The base model reproduced these dynamics only partially: simulated GMNN increased toward G2 in A-549, whereas CDT1 was non-monotonic, and in MDA-MB-468 both markers were largely flat across stages (left panels of Figures 4J-K, and Table S5). After fine-tuning *ProtiCelli* on 100 images per cell line (2,245 cells, corresponding to 4,490 marker-specific training images) for 8,000 iterations, the simulated intensities recovered the expected stage-dependent trends in both cell lines, with CDT1 declining and GMNN rising significantly from G1 to G2, matching the distributions measured in the real data (held-out test images: MDA-MB-468, 152; A-549, 222; middle versus right panels of Figures 4J-K, and Table S5).

Collectively, these results demonstrate that *ProtiCelli* can extract cell cycle-relevant information from morphological landmarks alone with minimal additional training, extending the model’s applicability beyond its original training domain. More broadly, this establishes a general fine-tuning framework in which users can generate targeted datasets to fine-tune *ProtiCelli* to specific cellular states or perturbations of interest.

### Unsupervised subcellular segmentation

Organelle segmentation in microscopy images typically requires specific organelle markers. As multiplexed IF beyond four markers is uncommon, most image analysis is limited to whole-cell or nucleus-cytosol segmentation, or to one organelle at a time [51–53]. Leveraging *ProtiCelli* ’s capability to generate images of organelle-specific proteins, we created a subcellular segmentation approach. We simulated 143 well-known organelle-specific proteins onto cell images (Table S6) and applied pixel-level spatial clustering [54] (see Methods; Figure S7A) to partition cells into 11 distinct subcellular regions, annotated by the most prevalent markers to identify subcellular regions. This revealed nuclear (nucleoli, nuclear speckles, nucleoplasm, nuclear membrane) and cytosolic compartments (mitochondria, ER) (Figure 5A). This approach extends to 3D by clustering across z-slices independently simulated (Figure 5B).

**Fig. 5:**
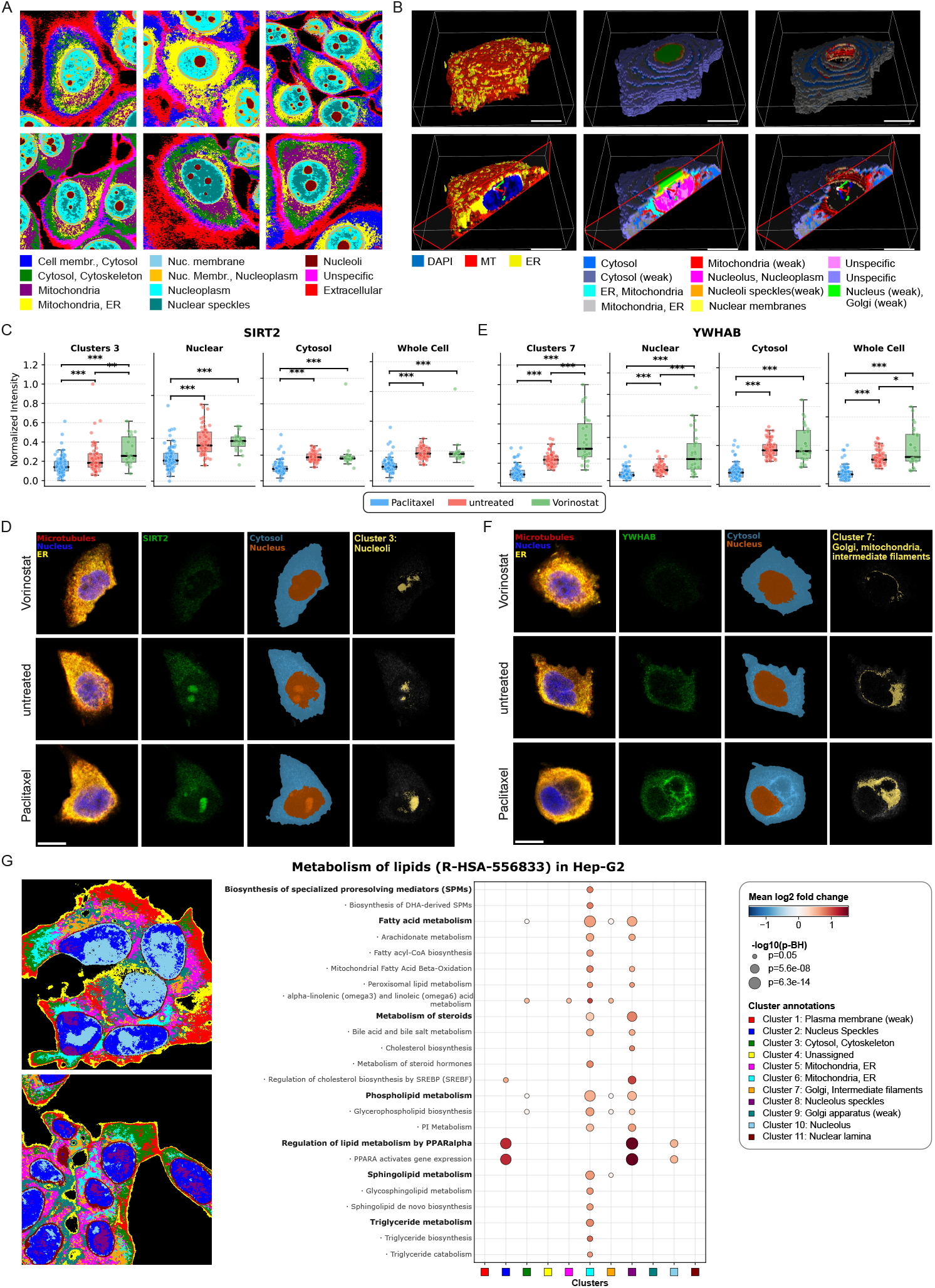
***ProtiCelli* enables unsupervised segmentation of subcellular compartments and proteome-scale spatial analysis and visualization applications.** (**A**) Representative spatial clustering of 143 organelle-marker proteins in MCF-7 cells. Pixel-level clustering of *ProtiCelli* -simulated images partitions cells into subcellular compartments; colors denote cluster identity. (**B**) Example views from *ProtiCelli* -based spatial clustering of an MDA-MB-468 cell applied to a 3D image stack, shown for selected clusters. Scale bar, 5 µm. (**C**) Normalized SIRT2 intensity across *ProtiCelli* -defined compartments (Cluster 3, nucleoli; nucleus; cytosol; whole cell) under Paclitaxel, untreated, and Vorinostat conditions; significance, Mann–Whitney *U* -test. (**D**) Representative single-cell images showing the cellular reference (microtubules, red; nucleus, blue; ER, yellow), the SIRT2 channel, nucleus and cytosol masks, and the Cluster 3 mask under each treatment condition. Scale bar, 10 µm. (**E**) Compartment-stratified YWHAB intensity in Cluster 7 (Golgi apparatus, mitochondria and intermediate filaments), nucleus, cytosol and whole cell. (**F**) Representative images of YWHAB and three *ProtiCelli* -defined regions, displayed as in **D**. Scale bar, 10 µm. (**G**) Spatial analysis of the Metabolism of Lipids gene set (Reactome, R-HSA556833) in Hep-G2 cells. Of the 758 proteins in the gene set, 566 were available for *ProtiCelli* simulation. *Left*, spatial cluster maps colored by cluster identity. Clusters were defined by pixel-level clustering of simulated organelle-marker images as in **A**. *Right*, spatial enrichment of descendant Reactome gene sets within each organellebased cluster. Binarized occupancy of each simulated protein image was aggregated within cluster masks and expressed relative to all other clusters (Methods). Dot color indicates the mean log_2_ spatial enrichment of detected proteins in each pathway– cluster pair; dot size indicates significance determined using a one-sided Wilcoxon signed-rank test. Only pairs with an FDR-adjusted *P <* 0.05 are shown. Pathways are arranged hierarchically on the *y* axis, with children of R-HSA-556833 in bold and grandchildren in plain text; spatial clusters are shown on the *x* axis.

*ProtiCelli* ’s subcellular segmentation enables unsupervised spatial quantification without dedicated organelle stains. In the *CM4AI-MDA* dataset, SIRT2 showed visually apparent nucleoli accumulation in Vorinostat-treated versus untreated cells, which whole-cell and nuclear measurements would miss but *ProtiCelli* -based segmentation resolved through segmenting isolated nucleolar regions (Figures 5C-D and S7B-C). Similarly, YWHAB showed perinuclear cytosolic accumulation under Vorinostat treatment, a localized change that subcellular segmentation revealed more precisely than whole-cytosol measurements (Figures 5E-F and S7D-E). This exemplified a broader framework for analyzing microscopy images with limited reference stains. While currently requiring microtubules and ER reference markers, the method enables systematic quantification of subcellular protein localization across fine subcellular compartments from a single standard IF experiment.

### Spatially disentangling gene sets

We next applied *ProtiCelli* ’s subcellular segmentation to systematically characterize the spatial organization of a gene set from *Proteome2Cell*. We segmented Hep-G2 liver cells into organellar regions and assessed the spatial organization of proteins involved in lipid metabolism (Figures 5G and S8). Nuclear regions were enriched for metabolic gene expression programs like ”PPARA activates gene expression”, whereas the multi-organellar region annotated with Mitochondria and ER was prominently enriched for ”fatty acid metabolism” and ”phospholipid metabolism”. ”Linoleic acid metabolism”, known to occur at the ER, extended to additional ER/cytosol clusters. Our virtual experiment showed that *ProtiCelli* and *Proteome2Cell* add a new datadriven spatial dimension to Reactome and Gene Ontology databases, disentangling terms and localizing where each process most likely occurs in a cell. Our method can also cluster directly on the gene set’s own proteins rather than organelles, revealing functional cellular regions more directly and with less organelle focus (Figure S7F).

### Spatially simulating protein assemblies

Beyond curated gene sets, *Proteome2Cell* can spatially resolve experimentally derived protein groups at a scale inaccessible to experiment. For example, we visualized 275 assemblies in U2OS cells that were recently identified by integrating microscopy and biophysical interaction data [55]. For each high-confidence assembly, we simulated protein assembly visualizations within individual cells, which recapitulated expected localizations (Figure 6A).

**Fig. 6:**
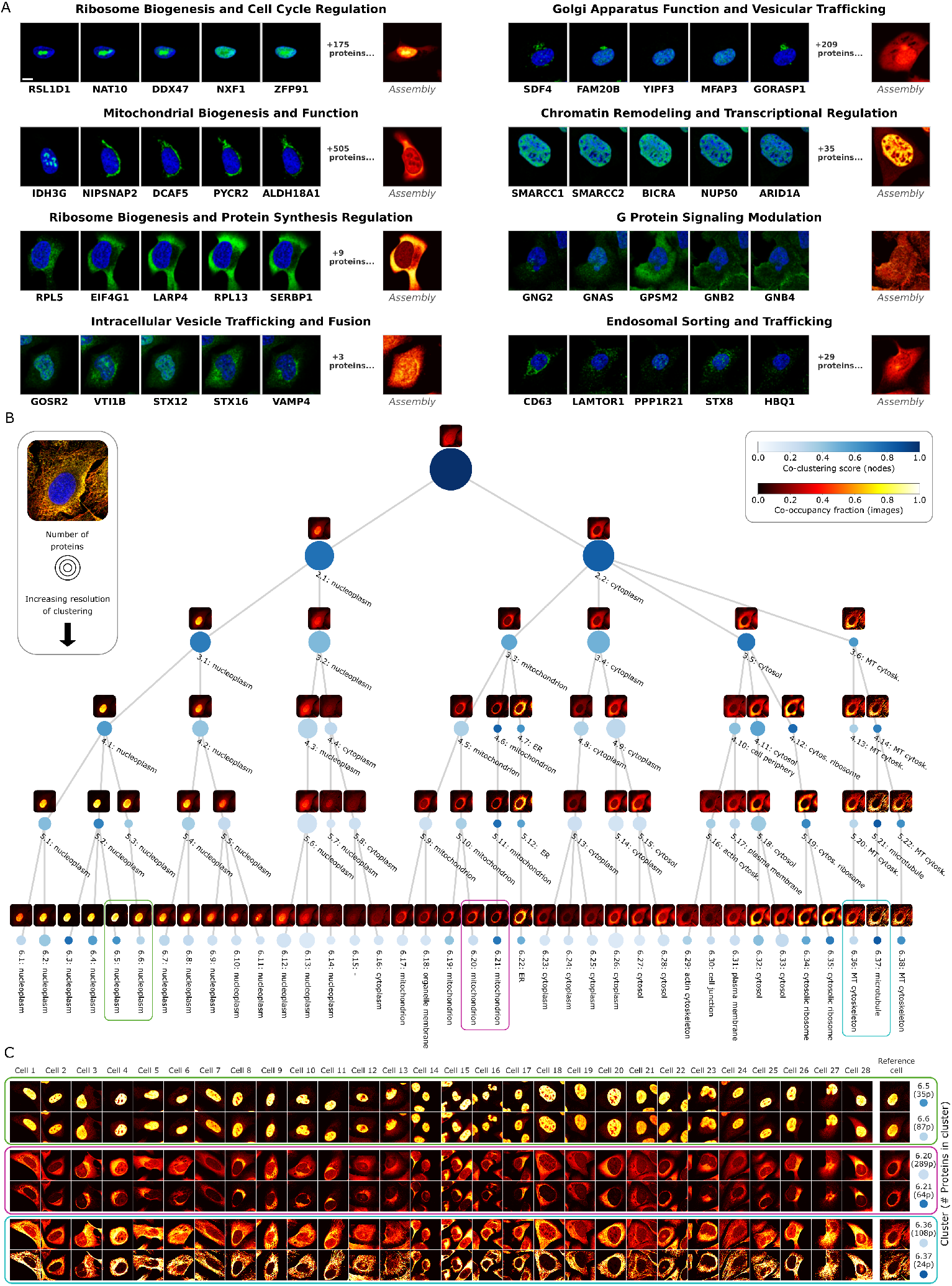
***ProtiCelli* enables proteome-scale spatial analysis, assembly visualization, and hierarchical subcellular mapping.** (**A**) *ProtiCelli* simulation of protein assemblies identified in [55]. Simulated immunofluorescence images of five representative constituent proteins shown alongside a heatmap of consensus expression across all assembly proteins. Scale bar, 10 µm. (**B**) Hierarchical multi-scale map of a single cell derived from a 12,800-plex *ProtiCelli* -simulated image. Each node represents a spatial cluster at one resolution; node color indicates the mean co-clustering score across 200 single-cell maps, defined as the fraction of cells in which protein pairs assigned to the same cluster in the reference cell co-clustered (Methods; top color bar). Heatmaps show the consensus expression of all proteins in each cluster in the reference cell; brightness indicates protein-expression overlap (bottom color bar). Colored frames denote clusters shown in **C**. , (**C**) Selected cluster heatmaps from the hierarchical map of the reference cell (**B**) compared with heatmaps for the same cluster proteins in other cells. The clusters shown correspond to pairs of highly and less stable clusters. Heatmaps for other clusters are shown in Fig. S9.

### Proteome-wide models of single cells

A contemporary goal of AI in biology is to build virtual cells [29]. *ProtiCelli* allowed us to generate 12,800-plex images of single cells (*Proteome2Cell* dataset), representing the first image-based virtual cell models of individual cells including nearly the whole cell proteome. To make these high-dimensional models interpretable, we generated single-cell multiscale maps for the 200 U2OS cells in the *Proteome2Cell* dataset. These allowed us to compare the single-cell specificity of subcellular protein architecture. By comparing the clusters across individual cells, we scored protein clusters by how conserved versus cell-specific they are (Figure 6B). Because these scores are derived from simulated images, we confirmed that they reflect real biology: clusters scoring more conserved occupied significantly tighter regions of the real-image SubCell single-cell embedding space, a correspondence that held across each evaluated hierarchy level (levels 4–7; Figure S10).

Conserved protein clusters represented key cell structures like the microtubule cytoskeleton, mitochondria, nucleoplasm, or ER (Figure 6B). More cell-specific clusters reflected dynamic processes like vesicular transport or cell adhesion. For the key cell structural markers, the maps distinguished variable clusters from the stable clusters (Figure 6C). The variable clusters mostly referred to proteins that multi-localize and show different subcellular localization across different cells, as revealed by cluster heatmaps.

We conclude that *ProtiCelli* -based virtual cell modeling enables unprecedented comparison of whole-proteome subcellular architecture across individual cells and annotates multi-scale cell models with information on cluster dynamicity, extending maps with additional functional information.

### A virtual cell in the Human Protein Atlas

To maximize utility by the research community, we have integrated a subset of *Proti- Celli* -simulated 12,800-plex image stacks into the HPA database (v26; to be released in Nov 2026). This adds over 1.4 million simulated images, extending the experimentally constrained protein-cell line coverage (Figure S11) to all proteins across all cell lines. An interactive virtual cell interface enables users to explore protein expression patterns, subcellular compartments, and complete signaling pathways within example cells, side by side with existing HPA experimental data to bridge computational predictions with the original experimental observations. The *ProtiCelli* -enhanced HPA marks a shift in spatial cell biology resources: from single-protein catalogs to proteome-scale virtual cells. This enables researchers to explore subcellular organization without specialized infrastructure or computational expertise by democratizing proteome-scale spatial analysis.

## Discussion

Capturing protein localization at subcellular resolution is essential for understanding both individual protein function and systems-level cellular organization. However, even advanced multiplexed imaging is limited to tens of markers, far from achieving comprehensive views of the thousands of proteins in a cell. *ProtiCelli* provides a computational framework for proteome-scale spatial analysis, generating high-resolution protein localization patterns for 12,800 proteins from basic cellular landmarks.

*ProtiCelli* outperforms other state-of-the-art models (*PUPS* [26] and *CELL-Diff* [27]) in pixel-level accuracy, morphological fidelity, and perceptual validity. Beyond technical metrics, we validate that *ProtiCelli* captures biologically meaningful hierarchical spatial organization down to protein co-localization, and can reflect context-dependent expression changes.

Exploring *ProtiCelli* ’s strengths, we developed new analytical approaches for spatial biology that broadly pave new avenues for microscopy-centered as well as general biology and proteomics research. We demonstrate that *ProtiCelli* spatially visualizes predicted protein assemblies, spatially disentangles gene ontology protein sets and signaling pathways, and segments subcellular compartments without specialized markers in bioimage analysis. Additionally, *ProtiCelli* supports systems-level analysis by enabling the construction of hierarchical spatial maps of subcellular organization in single cells, facilitating systematic comparisons across cell lines, cell states, and perturbation conditions.

Benchmarking and applying *ProtiCelli*, we noticed four key limitations that present opportunities for future improvement: First, *ProtiCelli* initially showed limited accuracy for cell cycle markers, but fine-tuning on a small dataset substantially improved performance. This demonstrates *ProtiCelli* ’s customizability, enabling lab-in-the-loop workflows where computational predictions guide experiments that refine the model for a specific task. This observation also demonstrates that *ProtiCelli* ’s limited capacity to recapitulate single-cell variability stems from a limitation in the number of cell states per protein in the training data and can be overcome by adding additional data. Second, *ProtiCelli* was trained exclusively on standard cell lines under normal conditions. Generalization tests on drug-treated MDA-MB-468 cells showed comparable performance to HPA, but incorporating diverse perturbed states during training could extend applicability to disease contexts and experimental perturbations. Third, *ProtiCelli* encodes proteins through learnable embeddings and cannot generalize to unseen proteins, a design choice that we made to focus on high fidelity. Finding a way to incorporate pretrained protein language model features without sacrificing prediction accuracy could enable predictions for uncharacterized proteins and cross-species transfer. Fourth, *ProtiCelli* ’s accuracy in positioning subcellular patterns in the cell depends on the structural information provided by the reference channels. Using additional reference markers could improve predictions for proteins whose localization is weakly constrained by the current nucleus/ER/microtubule combination.

*ProtiCelli* democratizes spatial proteomics by requiring only standard microscopy infrastructure and simple multiplexed IF images to achieve proteome-scale analytical capabilities. The integration of all simulated data into an interactive HPA interface democratizes proteome-scale spatial analysis, enabling researchers worldwide to explore models of cellular organization. Furthermore, *ProtiCelli* enabled predictive experimental design, in which computational hypotheses prioritize proteins for targeted validation, optimizing resource allocation while expanding the scope of biological discovery.

*ProtiCelli* establishes the feasibility of learning a generic generative model of proteome-wide spatial organization at single-cell resolution. By demonstrating robust generalization to unseen cell lines and broad applicability in spatial biology, *ProtiCelli* serves as a foundation model for computational spatial proteomics. Looking forward, *ProtiCelli* ’s framework could extend to disease modeling, drug mechanism studies, and systems-level investigation of spatial protein networks, potentially transforming how we understand and manipulate cellular organization in health and disease.

## Methods

### Experimental procedures

#### Generation of *FUCCI-CellCycle* dataset

The experimental workflow followed the HPA subcellular imaging protocol [7]. A-549 or MDA-MB-468 cells were seeded in fibronectin-coated glass-bottom 96-well plates and incubated for 24 hours. Cells were washed with PBS and fixed with ice-cold 4% paraformaldehyde for 15 min, followed by three washes with PBS. Cells were permeabilized with 0.1% Triton X-100 in PBS, through three consecutive incubations for 5 min. Next, cells were incubated overnight with primary antibodies dissolved in blocking solution (PBS + 4% FBS): mouse IgG1k antibody against Alpha Tubulin (Abcam, AB7291, RRID:AB 2241126, 1:500) to label microtubules, chicken antibody against calreticulin (Abcam, ab2908, RRID:AB 303403, manually conjugated to CF725 (Biotium, #92583) and thereafter used at 1:100) to label the ER, rabbit antibody against CDT1 conjugated to AlexaFluor647 (1:100, Abcam, AB211857-1002), rabbit antibody against GMNN conjugated to CL488 (1:200, Proteintech, CL48810802, RRID:AB 2918997). Next, cells were washed four times with PBS (10 min, RT). Cells were incubated with secondary antibodies for 90 min in blocking buffer: anti-mouse-IgG1k antibody coupled to AlexaFluor555 (1:2000, Invitrogen, A-21127). Next, cells were incubated for 10 min with DAPI solution in PBS. Next, cells were washed three times with PBS (10 min each) and imaged on a Leica Stellaris microscope equipped with a 63x water-immersion objective. Each channel was recorded as an independent series to avoid bleed-through. Scanning speed was 700 Hz, field of view size 2048x2048, resulting in a pixel size of 0.07 µm. 3D image stacks with an inter-plane distance of 0.7 µm were acquired.

#### Generation of 3D stack images for subcellular segmentation

MDA-MB-468 cells were seeded into 96-well plates and stained and imaged as described above for the *FUCCI-CellCycle* dataset except for different antibodies being applied. Primary antibodies: alpha-tubulin antibody (Abcam, RRID:AB 2241126, 1:1000) for labeling microtubules and a calreticulin antibody (Abcam, ab2908, RRID:AB 303403, 1:800) for labeling the endoplasmic reticulum. Secondary antibodies: Goat anti-Mouse IgG (H+L) Highly Cross-Adsorbed conjugated with Alexa Fluor 555 (1:800, ThermoFisher Scientific, A-21424, RRID:AB 141780), Goat anti-Rabbit IgG (H+L) Cross-Adsorbed Secondary Antibody conjugated with Alexa Fluor 488 (1:800, ThermoFisher Scientific, A-11008, RRID:AB 143165), and Goat anti-chicken IgY (H+L) conjugated to Alexa Fluor 647 (1:800, ThermoFisher Scientific, A-21449, RRID:AB 2535866).

### Image processing and data curation

#### HPA-training and HPA-testing datasets

Our dataset is built upon the Subcellular section of the HPA database (version 24), comprising images of stainings for 12,800 proteins (defined by unique Ensembl IDs) across 39 cell lines using 15,769 antibodies. Of these, 2,941 proteins were stained by multiple different antibodies; for these, images from multiple stainings were compiled together for model training. Some HPA antibodies bind multiple target proteins simultaneously (when the sequence similarity is very high), with resulting images representing all detectable proteins. For 462 such multi-targeting antibodies, targeted proteins were consolidated into single joint protein entries. In total, our dataset spans 13,659 proteins consolidated into 12,800 unique protein entries (single proteins or protein groups; Table S1). For each field-of-view (FOV) image, we applied intensity normalization (described below), downsampled to 0.75× the original resolution, and performed cell segmentation based on nuclear, microtubule, and ER markers. Individual cells were cropped using a 512×512 pixel window centered on the nuclear centroid. For cells near image boundaries, the window was shifted inward to ensure full coverage while retaining the cell. For each protein-cell line combination, we held out at least 3 single-cell images for testing and downstream analyses (e.g., *Proteome2Cell* assembly), with remaining images used for model training.

We normalized the pixel intensity range of different single-cell cropped images to ensure stable training dynamics as follows. Given a multi-channel image **I** ∈ R*^H^*^×*W*^ ^×*C*^ where *C* = 4 represents three cellular markers and one protein of interest, we clipped each image channel at its channel-specific 99.5th percentile and then normalized all channels using the post-clipping maximum intensity of the nucleus (DAPI) channel as a shared scaling factor:

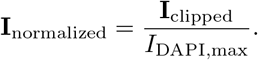

This shared normalization preserves relative intensity differences between channels; consequently, normalized endoplasmic reticulum (ER) and microtubule (MT) intensities may exceed 1. Finally, intensities were centered and scaled using

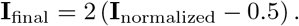

#### CM4AI-MDA dataset

Our dataset is built upon our CM4AI immunofluorescence dataset for MDA-MB468 cells (Release: Beta, June 2025 [42, 56]). The dataset comprises high-resolution confocal images of cultured MDA-MB-468 cells under three conditions: treated with Paclitaxel (NSC 125935, Selleckchem S1150), treated with Vorinostat (SAHA, Selleckchem S1047), or untreated.

The CM4AI dataset [56] was generated following the HPA subcellular imaging protocol [7], involving reference marker staining with DAPI for labeling nucleus, an alpha-tubulin antibody (Abcam, RRID:AB 2241126) for labeling microtubules, and a calreticulin antibody (Abcam, ab2908, RRID:AB 303403) for labeling the endoplasmic reticulum. The only difference from the HPA subcellular imaging protocol [7] is that the imaging was done on a Leica Sp8 Confocal Microscope. Microscope settings: Pinhole 1 Airy unit, 16-bit acquisition, 1.28x zoom, 2048x2048 pixels field of view, 700 Hz scan speed, resulting in a pixel size of 0.07 µm. All channels were detected on HyD detectors in multiple individual series to reduce bleed-through: (1) Nuclei (DAPI) and Endoplasmic Reticulum (Alexa 647), (2) the protein of interest channel (Alexa 488), (3) the microtubule channel (Alexa 555). The detector gain and the laser intensity for the protein of interest (Alexa 488) channel were constant for all images acquired. On every plate, a negative control (no primary rabbit antibody) was included and recorded to confirm that no background fluorescence signals were visible in the Alexa 488 channel.

The images in the CM4AI dataset [56] are in jpg format and already normalized (based on the tool: https://github.com/CellProfiling/TifCs To HPA-PNG-JPEG/) individually for each antibody staining triple (all three conditions) as follows: the upper 0.01 percentile is determined across all images and adjusted to 10,000, if it is less than 10,000; all images of that protein were re-scaled so that in the resulting jpg images, the maximum intensity (255) corresponds to the upper 0.01 percentile or 10,000 in the original images, whichever is higher. This normalization is applied for all channels individually.

To assemble the *CM4AI-MDA* dataset, we fetched the normalized JPG images for 453 proteins from the release (protein names are available in Table S2). We segmented and cropped the images to 1024x1024 single-cell-centered images using the https://github.com/CellProfiling/HPACellSegmentatorPortable tool, yielding 112,410 cropped images. Next, we normalized the images as described for the *HPA-training* and *HPA-testing* datasets. Next, for each perturbation set (Paclitaxel, Vorinostat, Untreated), we generated *ProtiCelli* images for all 453 proteins, measured average pattern intensities (segmented by 1.5× Otsu threshold) as a proxy for expression levels (see below), and compared predicted versus experimental expression changes (see Figures 4B and C for examples).

#### FUCCI-CellCycle dataset

The 3D confocal image stacks of A-549 and MDA-MB-468 cells contained five channels: DAPI, GMNN, ER, microtubules, and CDT1. Cell segmentation was performed on the DAPI channel in 3D using Gaussian smoothing (*σ* = 1), Otsu-thresholding, morphological closing, and per-slice binary hole-filling. The optimal z-plane for each cell was selected as the slice closest to the intensity-weighted nucleus centroid. Individual cells were cropped using a window determined by the nucleus bounding box at 512 512 pixels. For cells near image boundaries, the window was shifted inward to ensure full coverage while retaining the cell. Cells with a nuclear area smaller than 5,000 pixels or for which more than one quarter of the intended crop area fell outside the image boundary were excluded. This resulted in 3,092 and 1,896 single-cell image crops for A-549 and MDA-MB-468, respectively. Intensity normalization was applied per field of view for the DAPI, ER, and MT channels using the 99.9th percentile of pixel values, while GMNN and CDT1 channels were normalized globally using the 99.99th percentile computed across all fields.

#### Staining intensity calculations

When quantifying protein expression levels, we segmented protein expression regions using an adaptive threshold of 1.25× the Otsu threshold value, unless otherwise specified. Mean intensity was calculated as the average pixel intensity within the segmented region (i.e., pixels above the threshold). This procedure was applied to both real and simulated images, with pixel intensities in the range [0, 255].

#### Machine learning

*ProtiCelli* is adopted from a diffusion model, which has become the leading approach to high-fidelity image generation [30, 31]: it learns the full conditional distribution, produces sharp and diverse samples, and trains stably, avoiding model collapse and optimization instability. We implemented *ProtiCelli* as a conditional latent diffusion model [32] (Figure S1), which uses a pretrained Variational Autoencoder (VAE) (Stable Diffusion 3.5 [33]) for computational efficiency. The diffusion component is built on the Diffusion Transformer Large (DiT-L) architecture [34], We chose this transformerbased architecture over the convolutional U-Net because self-attention better captures long-range spatial dependencies and scales more effectively to large, diverse training data [35, 36]. For time-efficient proteome-scale image generation, we implemented the Elucidating Diffusion Models (EDM) framework [37].

#### Elucidating Diffusion Models

We employ the EDM framework [37] to train our generative model and sample from it. Following the EDM formulation, we precondition the data using *c*_skip_ 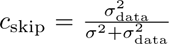, c_out_ 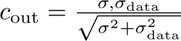, and *c*_in_ = 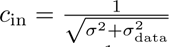, with *σ*_data_ = 0.5. The network is conditioned on the noise level using *c*_noise_ = ^1^ log *σ*.

Noise levels follow the Karras schedule

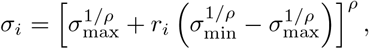

where *σ*_min_ = 0.002, *σ*_max_ = 80, and *ρ* = 7. For the training noise grid, *r_i_* = *i/N* for *i* = 0*, . . . , N* , with *N* = 1000; during training, indices were sampled uniformly from *i* = 0*, . . . , N −* 1. For inference, we used *T* = 50 denoising steps with *r_i_* = *i/*(*T −* 1) for *i* = 0*, . . . , T −* 1, followed by a final transition to *σ* = 0.

The generative model *g* receives the preconditioned noisy input *c*_in_*x*_noisy_, and its output is converted to a denoised estimate as

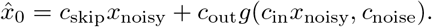

During inference, we use 50 Euler denoising steps.

#### Latent diffusion

*ProtiCelli* employs latent diffusion for efficient and scalable image generation. A pretrained variational autoencoder (VAE) from Stable Diffusion 3.5 [33] encodes both cellular landmark images and target protein images into 16 64 64 latent representations. Single-channel protein images are converted to three-channel RGB format via channel replication before encoding. These latent representations are processed by the diffusion model (described below) and decoded back to pixel space via the VAE decoder. Latent values (range ≈ -4 to 4) are scaled by 0.25 to normalize them to [ −1, 1] for diffusion training.

#### Neural network design

Our deep generative model follows a Diffusion Transformer Large architecture with a patch size of two (DiT-L/2). We adapted the standard DiT design to condition image generation on three cellular landmark channels—nucleus, endoplasmic reticulum (ER), and microtubules—as well as on target protein identity and cell-line identity.

The variational autoencoder jointly encodes the cellular landmark channels into a landmark latent **c** R*^B^*^×*C*cond×*H*×*W*^ , where *C*_cond_ = 16 and *H* = *W* = 64. At diffusion step *t*, the noisy target-protein latent **x***_t_* R*^B^*^×*C*out×*H*×*W*^ , where *C*_out_ = 16, is concatenated channel-wise with the landmark latent:

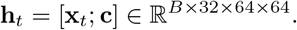

The combined latent is divided into non-overlapping *P × P* patches and linearly projected to token representations **z**_0_ *∈* R*^B^*^×*N*×*D*^, where

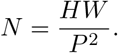

With *H* = *W* = 64 and *P* = 2, the model processes a 32×32 token grid (*N* = 1,024 tokens). The token embedding dimension is *D* = 1,024.

Protein and cell-line identities are embedded separately together with the diffusion timestep. The resulting timestep-conditioned protein and cell-line embeddings are

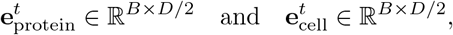

respectively. Because *D* = 1,024, each embedding has dimension 512. The two embeddings are concatenated to form the complete conditioning vector

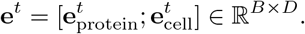

To enable cell-line-agnostic generation, cell-line conditioning is dropped stochastically during training: with probability 0.2, the cell-line identity is replaced by a learned null or default label. The same default label is used when cell-line information is unavailable during inference.

Each transformer block uses adaptive layer normalization with zero-initialized modulation (adaLN-Zero) to condition both the self-attention and feed-forward sublayers. The conditioning vector is passed through a SiLU activation and a linear projection:

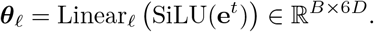

The projected vector is divided into shift, scale, and gate parameters for the selfattention and feed-forward sublayers:

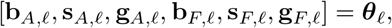

where each parameter lies in R*^B^*^×*D*^ and is broadcast across the token dimension.

For the self-attention sublayer, the normalized and modulated representation is

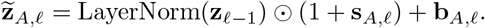

The attention output is gated before being added through the residual connection:

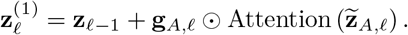

The feed-forward sublayer is modulated analogously:

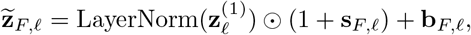

followed by

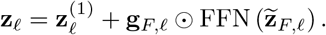

Here, ⊙ denotes element-wise multiplication and *ℓ* indexes the transformer block. The feed-forward network uses a GELU activation, expands the representation to an intermediate dimension of 4*D*, and projects it back to dimension *D*.

After processing through *L* = 24 transformer blocks, the final token representations undergo conditioning-dependent shift and scale modulation:

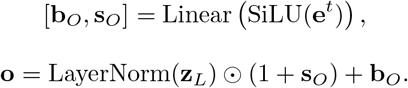

Each output token is then linearly projected to *P* ^2^*C*_out_ = 64 values:

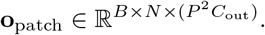

The projected patches are rearranged into the spatial latent representation

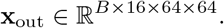

The model comprises 24 transformer blocks with 16 self-attention heads and a head dimension of 64, giving a total token embedding dimension of *D* = 16 × 64 = 1,024. It processes 1,024 tokens obtained from a 64 × 64 latent using 2 × 2 patches. The biological conditioning vocabulary covers 12,800 protein entries and 39 cell lines.

#### Model training

Training employed the AdamW optimizer (*β*_1_ = 0.9, *β*_2_ = 0.999) with learning rate 10^−4^ and weight decay 0.01. The learning rate followed a linear warm-up schedule over the first 5,000 iterations, increasing from 10^−7^ to 10^−4^. We maintained an exponential moving average (EMA) of model weights with decay parameter 0.9995 for evaluation. The model was trained for one million iterations on 8 NVIDIA H100 GPUs with a per-GPU batch size of 32 (total effective batch size 256).

#### Reliability-aware prediction by ensemble medoid selection

For each cell, the reference channels were encoded once into the latent space of the variational autoencoder and held fixed, after which *N* independent latents were drawn from the conditional diffusion model under a deterministic sampler, so that variability among ensemble members arose solely from the initial latent noise rather than from the conditioning. Each latent was decoded to a single-channel protein image *x*^*_i_* (*i* = 1*, . . . , N* ), and a cell region Ω was defined by thresholding the cellular reference channels. Pairwise agreement was quantified by the PCC *ρ*(*x*^*_i_, x*^*_j_*) evaluated over Ω, computed after a light Gaussian smoothing that attenuated high-frequency sampling texture without altering the underlying spatial pattern. Rather than reporting an ensemble mean, which blurs spatially incoherent structures such as vesicular puncta and yields an image that no individual draw resembles, the medoid

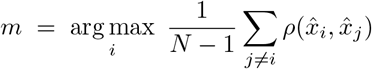

was retained as the emitted prediction *x*^*_m_*, preserving the high fidelity of a generated sample. Selection of the representative image and quantification of reliability were treated as separate operations: because the medoid centrality is a maximum over *N* noisy per-member agreements and is therefore biased upward by selection, increasingly so at small *N* , reliability was graded not on it but on the mean pairwise correlation of the whole ensemble,

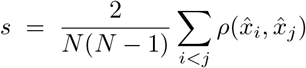

the lower-variance, selection-free estimator of ensemble coherence. Members that were non-finite or spatially constant within Ω were excluded from both the medoid selection and the score. The score *s* was reported as a continuous scalar; because it derives from correlation, it is invariant to per-image scale and offset and therefore comparable across proteins of differing expression level, and *N* was held fixed across all predictions so that scores remained directly comparable (*N* = 20 in this work). A higher value of *s* indicates that independent generations under fixed conditioning converged on a consistent localization, providing a per-prediction measure of the degree to which the reference channels constrained the inferred protein pattern.

### Model evaluation

#### Benchmarking methods

We evaluated *PUPS* [26] and *CELL-Diff* [27] within their native pipelines, using the officially released code and the preprocessing specified in each publication without modification or re-tuning, so that each model was applied as its authors intended. Protein sequences were filtered to reviewed UniProt entries with single isoforms and encoded using ESM2 embeddings following *PUPS* protocols.

Cellular landmark images (nucleus, ER, microtubules) were resized to match each model’s requirements: 3 *×* 128 *×* 128 (*C × H × W* ) for *PUPS* and 3 *×* 256 *×* 256 for *CELL-Diff*, determined by their respective architectures. These resolutions reflect the native architectures and released inference pipelines of the respective models. Images were normalized to [0, 1] for model input. All model predictions (*ProtiCelli*, *PUPS*, *CELL-Diff* ) were rescaled to [0, 255] to match ground truth images for quantitative evaluation.

Inference used a batch size of 32 for all models. *PUPS* used default sampling parameters, *CELL-Diff* used 200 diffusion timesteps with maximum sequence length 2048, and *ProtiCelli* used 50 EDM sampling steps.

#### Reference-channel determinability

For each protein, we quantified how much of its within-cell spatial organization is recoverable from the three reference stains — microtubules, nucleus and endoplasmic reticulum (ER) — with an information-theoretic score *X* computed on real HPA single-cell images (12,403 proteins; 31,810 protein–cell-line pairs across 12 cell lines; 1.04 million cells). Single cells were delineated by a four-model consensus segmentation (CellSAM, Cellpose-SAM, a Cellpose/DINO model and HPACellSegmentator, merged by probabilistic STAPLE) and analyzed at 512 × 512 within the target cell’s mask. Within each cell, the protein channel (after subtracting its in-mask median and clipping at zero) defines a normalized spatial mass *q* over in-mask pixels, and *L* = *D*_KL_(*q ‖* uniform) = log *N − H*(*q*) (bits; *N* = in-mask pixels) measures how spatially structured the protein is. From the reference channels we built a 21dimensional per-pixel texture descriptor (per channel: raw intensity, Gaussian blurs at *σ* = 2, 4, 8, 16 px, and the differences *G*_8_ − *G*_2_ and *G*_16_ − *G*_4_), rank-normalized within the mask, and partitioned it into texture states with a single, protein-blind MiniBatch *k*-means codebook. The joint descriptor is quantized at *K* = 96 and each single-channel descriptor at *K* = 32: the joint sees 21 features against a single channel’s 7, so equal *K* would force the joint to tile a threefold larger space with the same number of centroids and could place it below a channel whose context it strictly contains. Scaling *K* linearly in feature dimension is a lower bound on the correction rather than a tuned choice. The part of *L* explained by this reference context, *G*, was estimated outof-sample by leave-one-cell-out: enrichment learned from a protein’s remaining cells (shrunk toward no enrichment) predicts a per-pixel distribution *p* for the held-out cell, giving *G* = max(0*, L D*_KL_(*q ‖ p*)). Determinability is *X* = *G/ L*, summed over a protein’s cells before taking the ratio, so *X* ∈ [0, 1] is the fraction of within-cell positional information recoverable from the reference channels. Because a finite codebook cannot perfectly reconstruct a continuous channel, a reference channel supplied as the protein scores below 1 (0.72, 0.89 and 0.74 for microtubules, nucleus and ER; mean 0.78); we therefore report *X* divided by this ceiling, capped at 1.

The spatial relationship between a protein and reference channels is rarely simple co-localization. Proteins may surround, be surrounded by, or sit adjacent to landmark stains, e.g., nucleoli occupying DAPI-dark regions within an otherwise bright nucleus. Standard same-pixel co-localization scores miss these complex relationships entirely.

While complex image models can capture these spatial dynamics, using a trained network to evaluate *ProtiCelli* would introduce circularity through shared inductive biases. To avoid this, we prioritized validity over complexity. We define our determinability metric (*X*) using a fixed, protein-blind vocabulary of local landmark context (multi-scale filters quantized into *K* states), scored by a held-out statistical information measure.

Three properties should be considered when interpreting *X*:

- **Robustness.** *X* is not a tuning artifact. Varying the joint codebook over *K* = 32– 1024 leaves protein rankings essentially unchanged (Spearman *ρ* ≥ 0.99 against *K* = 32), and the absolute scale varies little over that range (compartment-median *X* of 0.239, 0.251, 0.254 and 0.249 at *K* = 32, 96, 256 and 1024). Only at *K* = 4096 does *X* fall below its *K* = 32 value (0.225), where too many texture states are being estimated from the cells of a single protein; the reported values therefore sit on the stable part of this range rather than at an extreme of it.
- **Reproducibility.** By leaving out whole cells to estimate held-out information, *X* captures only structural mappings that recur across cells. Consistent localizations, like microtubule-associated proteins aligning with the cytoskeletal network, score well. Conversely, reference-linked but cell-idiosyncratic placements, such as dynamically positioned endosomes, score low.
- **Instability for diffuse proteins.** For proteins lacking distinct spatial structure, such as broadly diffuse cytosolic proteins, the mass is near-uniform (small *L*). This makes the ratio *X* = *G/ L* computationally unstable. In this regime, a low *X* reflects a weak positional signal to explain, rather than a confident measurement of low determinability.

Together, these properties make *X* a deliberately conservative estimator. The determinability of dynamically positioned or broadly diffuse localizations may therefore be underestimated, so a low *X* in these cases should be read as a lower bound rather than as evidence of genuinely random placement.

#### Protein embedding generation and preprocessing

Protein embeddings were generated using *SubCell* [43], a microscopy foundation model that captures single-cell patterns of subcellular protein localization and cell morphology. Specifically, we employed the MAE-CellS-ProtS-Pool variant. For each protein, four-channel IF images (nucleus, ER, microtubules, and protein of interest) were processed through the encoder to obtain 1,536-dimensional embedding vectors.

##### Embeddings of real images

Reference embeddings were derived by embedding all crops from the training and test sets combined. Cells were then filtered and aggregated as described in [43]: cells lacking a subcellular location annotation were discarded, as were antibodies mapping to more than one gene and antibodies of Uncertain gene reliability; for each protein a single best antibody was retained by reliability rank (Enhanced *>* Supported *>* Approved). Protein-level representations were obtained by averaging the surviving single-cell embeddings for each protein–cell-line combination, and the location field of an aggregated protein is the union of its pooled annotations.

##### Embeddings of simulated images

For each superplexed cell from *Proteome2Cell*, each protein channel was combined with the corresponding three cellular landmark channels (nucleus, ER, microtubules) and processed through *SubCell*.

##### Depth matching

The analyses that compare real against simulated data — the joint projections, the label transfer, and the hierarchical clustering comparison — additionally depth-match the two sides; analyses on real data alone do not. A protein has far more simulated than real cells available (median 20 real cells per U2OS protein against 200 simulated), so averaging all available cells would give the simulated side a systematically less noisy protein mean. For every protein we therefore drew, without replacement, exactly as many simulated cells as that protein has real cells, and averaged those. Depth matching was repeated at three independent sampling seeds; the real side is unaffected by construction, since only the simulated draw is reseeded.

##### Embedding alignment

Every comparison is restricted to the proteins present in both compared datasets, with protein ordering synchronized so that rows correspond between matrices; for the real-versus-simulated comparisons this intersection comprises 27,399 protein–cell-line pairs across the twelve lines. Each arm was then centered by subtracting its centroid vector to remove the constant offset between the two arms, which otherwise dominates any joint embedding and produces an apparent real-versus-simulated separation that is an artifact of the arms’ differing output ranges rather than of localization content.

#### Dimensionality reduction and visualization

UMAP [57] was used to visualize how the embeddings are distributed. For each cell line, the depth-matched real and simulated protein means were projected jointly so that both arms share one set of coordinates. We used a *k*-nearest neighbor graph with *k* = 125 neighbors computed on the full 1,536-dimensional embeddings, Euclidean distance and *min dist* = 0.1. Maps were colored by data source (real vs. simulated) or by dominant subcellular location. Because each cell line is projected independently and a UMAP map has no canonical orientation, every panel was rigidly rotated and reflected onto the largest cell line by orthogonal Procrustes on the proteins the two share, without scaling. This is an isometry, so it leaves all reported statistics unchanged and serves only to give the panels a common orientation. Coordinates remain comparable only within a panel.

Points were colored by each protein’s main HPA location, taken as the first-listed annotation for that protein and cell line and applied to both arms alike, so that any label noise affects the real and simulated panels identically. Most annotated proteins carry more than one location, and giving multi-localizing proteins their own category would place the majority of the cloud in a single bucket; using the main location instead keeps the coloring interpretable. For visualization clarity, fine-grained HPA location annotations were consolidated into broader categories: nucleolar substructures (nucleoli, nucleoli fibrillar center, nucleoli rim) were merged into ”Nucleoli”; cytoskeletal components (microtubules, actin filaments, intermediate filaments) were merged into ”Cytoskeleton”; and endomembrane system components (ER, Golgi apparatus, vesicles) were merged into ”Endomembrane system.”

We calculated two summary statistics to evaluate the embedding spaces. For both metrics, we identified the 15 nearest neighbors for each protein using the full 1,536-dimensional embeddings rather than the two-dimensional projections. First, to quantify *arm separation* (the extent to which real and simulated protein means occupy distinguishable regions), we computed the mean fraction of a protein’s neighbors that originated from the same arm. In this context, a value of 0.5 represents random chance for a two-arm mixture of equal size. Second, to determine *location purity*, we evaluated a single arm and calculated the mean fraction of a protein’s neighbors that shared its annotated subcellular compartment. We display both metrics on each panel and tabulate them per cell line alongside the figure.

#### Cross-modal label transfer

To ask whether embeddings of simulated images carry the localization information needed to annotate a protein, we transferred subcellular location labels between arms by *k*-nearest neighbors (*k* = 5, cosine distance) under a one-index design: one arm supplies the labeled index and the other the queries. Performance is reported as top1 accuracy and macro-averaged F1 over location classes, cross-validated in five folds grouped by protein so that no protein appears in both index and query, and averaged over the three depth-matching seeds. To bound our interpretations, we created a morphology-only baseline by blanking the protein channel prior to SubCell encoding. This isolated the cell morphology signal contributed by the landmark stains, enabling us to measure the predictive power of the reference channels alone.

#### Hierarchical map of the human proteome

To quantitatively assess how faithfully the generative model reproduces the hierarchical protein localization structure present in real HPA data, we compared Leiden clustering labels across seven hierarchy levels. Clustering was performed independently on protein-level *SubCell* embeddings of real (HPA) and simulated images in U2OS cells using the procedure described in [43] using Python 3.10 and ScanPy 1.10.2. Using the 9,533 proteins shared by the real and depth-matched simulated U2OS data, we performed iterative subclustering with the Leiden algorithm [58]. Resolution was increased while neighbor counts were decreased (125 to 10) to move from global to local structure, giving 1, 2, 6, 14, 22, 38, and 86 clusters at hierarchy levels 1–7. This creates a hierarchical graph in which edges represent the derivation of clusters across successive resolutions. To enable a direct comparison, the resolution for the simulated data was tuned so that each level yields the same number of clusters as the corresponding real level.

Matching the cluster count alone does not determine a partition: at level 2 a resolution that separates a small group of proteins from the rest and one that splits the proteome near-evenly both yield two clusters. Among all resolutions attaining the target count, we therefore select the one of maximum modularity, which distinguishes these two outcomes without imposing any explicit cluster-size threshold; where the resolution grid contains no target-attaining value, the resolution closest to the target is used. Leiden optimization is run to convergence.

Because Leiden and the neighbor graph depend on a random seed, the full ladder was run at 30 seed combinations (three depth-matching seeds × ten clustering seeds), and all clustering results are reported as the mean ± one standard deviation across those runs.

Leiden cluster identifiers are arbitrary labels. For presentation, clusters were renumbered 0, 1, 2*, . . .* from left to right at each level of the drawn tree. This is a relabeling only: the partitions are unchanged and the agreement metric below is invariant to it.

#### Functional enrichment

Clusters were annotated using g:Profiler v1.0.0 [59] against the human genome. We applied GO Cellular Component for levels 1–6 and GO Biological Process/Molecular Function for level 7. The g:SCS algorithm provided multiple testing corrections, and a significance threshold of *p <* 0.05 was used to label each node with its top enriched term.

#### Quantitative clustering comparison

At each hierarchy level, the agreement between the two sets of clustering labels was quantified by the Adjusted Mutual Information (AMI) [60]. AMI is corrected for chance agreement and normalized to lie in [0, 1], with 1 indicating identical partitions. Agreement was computed at all 30 seed combinations and is reported as mean ± one standard deviation.

An absolute AMI is difficult to read without knowing how much agreement is attainable: even two independent clusterings of the *same* data disagree, because a protein mean estimated from a finite number of cells is noisy. We therefore estimated a per-level ceiling by splitting the real cells of each protein into two disjoint halves, aggregating each half separately, and clustering the two halves through the identical pipeline. The resulting real-versus-real agreement is the highest AMI the protocol could return at that level given measurement noise, and the real-versus-simulated value is interpreted against it. Both series were computed over the same 30 seed combinations so that the two are directly comparable. To keep the ceiling a like-for-like bound, the split-half arms use one antibody per protein, matching the main analysis.

#### Hierarchy tree rendering

The real and simulated hierarchies are each drawn as a tree in which nodes are clusters, node area is proportional to the number of member proteins, and an edge joins a cluster at one level to a cluster at the next when their protein sets overlap. Node color encodes the significance of the top enriched term for that cluster, and each node is captioned with its level, its cluster identifier, and that term. Trees are laid out topdown with graphviz. Because the two arms are clustered independently, one tree is drawn per arm, and no correspondence between their cluster identifiers is implied. A single representative seed combination is shown.

#### Subcluster visualization

To show *where* the two clusterings agree or diverge, hierarchy levels 2-4 are displayed as a two-by-two panel: rows are the real and simulated data, columns are the clustering derived from the real and from the simulated arm, so the same points are recolored by each partition in turn. Each arm is projected separately here (same neighbor count, metric, and seed as above), and the simulated map is rotated, reflected and translated onto the real one by orthogonal Procrustes so the two rows correspond shape for shape; cluster identifiers are matched between columns by maximum overlap so that a color means the same group across panels. Because the two arms are projected independently, their coordinate scales are arbitrary and unrelated; the simulated map was therefore rescaled along each axis so that both arms span the same width and height. These are presentation transforms on the two-dimensional maps only: the clusterings themselves are computed on the 1,536-dimensional embeddings, and no reported number depends on the map coordinates.

#### Consensus co-clustering stability of single-cell hierarchy trees

To assess whether the protein clusters identified in a given simulated cell reflect reproducible co-localization patterns rather than cell-specific noise, we colored each cell’s hierarchy tree by the consensus co-clustering frequency computed across all 200 simulated cells.

#### Per-cell hierarchical clustering

For each simulated cell, *SubCell* embeddings were obtained for all proteins present in that cell. Leiden clustering was applied iteratively at increasing resolution to construct a seven-level hierarchy matching the target cluster counts used for the U2OS embeddings of real images (1, 2, 6, 14, 22, 38, and 86 clusters at levels 1–7, respectively). At each level, the Leiden resolution parameter was tuned independently for the simulated cell embeddings to reproduce the target cluster count. The resulting per-cell cluster label table records, for each protein, its cluster assignment at every hierarchy level.

#### Consensus co-clustering matrices

Independently of the hierarchical clustering, flat Leiden clustering was run on each of the 200 superplexed U2OS cells separately at six resolutions *γ∈ {*0.10, 0.25, 0.50, 0.75, 1.00, 1.50*}*. For each cell *c* and resolution *γ*, a binary cooccurrence matrix *A*^(*c,γ*)^ was constructed, where 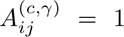 if proteins *i* and *j*were assigned to the same Leiden cluster and 0 otherwise. The consensus matrix at resolution *γ* is the cell-averaged co-occurrence frequency:

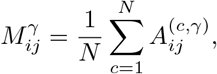

where *N* = 200 is the total number of cells. Each entry *M ^γ^ ∈* [0, 1] thus represents the fraction of cells in which proteins *i* and *j* were co-clustered at resolution *γ*.

#### Cluster cohesion scoring

Each hierarchy level *ℓ* 2*, . . . ,* 7 was paired with the consensus matrix at the corresponding resolution (levels 2–7 map positionally to *γ* = 0.10, 0.25, 0.50, 0.75, 1.00, 1.50, respectively; the root level 1 is excluded). For each cluster *k* at level *ℓ* containing a set of proteins *_k_*, the *cohesion* was defined as the mean pairwise consensus co-clustering frequency among cluster members:

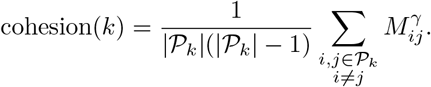

Single-protein clusters receive a cohesion of 1. High cohesion indicates that the proteins in the cluster are consistently co-clustered across many cells; low cohesion indicates cell-specific or unstable groupings.

#### Cluster-stability validation

To test whether these model-derived cohesion scores reflect real biology, we compared them against the spread of each cluster’s proteins in the SubCell embedding space of *real* HPA images (Figure S10). For each cluster, we pooled the real U2OS singlecell SubCell embeddings of all member proteins with at least 10 real cells (295,893 U2OS cells in total; clusters with fewer than 50 pooled cells were excluded) and measured cloud spread as the mean pairwise cosine distance. To ensure the relationship was not a cluster-size artifact, spread was computed at a fixed protein budget by bootstrap (*B* = 100 resamples, each drawing *p*_sub_ member proteins and at most 20 cells per protein; the reported spread is the resample mean and the 2.5th/97.5th percentiles give each cluster’s 95% confidence interval). The budget was set per level to the smallest retained cluster’s protein count, capped at 200, so that no cluster was dropped (*p*_sub_ = 34, 22, 22, 6 at levels 4–7). At each level we correlated cohesion against real-cloud spread using Spearman’s *ρ*(primary, with Pearson’s *r* reported alongside), testing each level independently rather than pooling across the tree hierarchy. We report levels 4–7 (14–86 clusters); coarser levels are omitted because each of their clusters spans thousands of proteins or whole compartments rather than coherent co-localization groups. Embeddings of simulated images were used only to define clusters, never to measure spread. Finally, to confirm the internal consistency of the model’s two Figure 6B scores, we correlated the co-clustering (cohesion) score with the co-occupancy score across the 38 leaf clusters using Spearman’s *ρ* (primary, with Pearson’s *r* reported alongside).

#### Tree visualization

The hierarchy tree for the single cell was constructed using the same iterative subclustering procedure described above, applied to that cell’s protein embeddings. Each node is colored according to its consensus cohesion score using a yellow–orange–red colormap (low to high cohesion). Node area is proportional to cluster size, and clusters with fewer than three proteins are omitted for clarity. The resulting figure shows one representative tree from a single simulated cell (cell 134), illustrating how consistently each protein grouping is recapitulated across the full population of 200 simulated cells.

#### Perturbation response quantification and virtual profiling

For each of the 453 proteins with sufficient data, we applied Mann–Whitney U tests to assess whether single-cell expression distributions differed significantly (*p <* 0.05) between treated and untreated conditions, separately for real and simulated images. Each protein was thereby classified into one of three states: non-significant, significantly upregulated, or significantly downregulated, based on the sign of the median expression difference between conditions. A protein was scored as a correct match if the simulated images exhibited both statistical significance and a concordant direction with the real images. We defined two complementary evaluation metrics: precision (correct matches divided by the total number of proteins predicted significant in the simulated images) and recovery rate (correct matches divided by the total number of proteins significant in the real images). To test whether the observed agreement exceeded chance, we performed permutation tests with 100,000 iterations. Under the null hypothesis that the simulated images carry no information about drug-specific perturbation effects, we randomly reassigned each protein’s real three-state outcome among all tested proteins while holding the *ProtiCelli* predictions fixed, and computed the number of correct matches in each permutation. Because precision and recovery are both functions of the same count of correct matches, they share a single permutation p-value. This null model accounts for the marginal distributions of both significance prevalence and directional bias, providing a conservative baseline that reflects the high fraction of proteins affected by each drug. P-values were computed as the fraction of permutations yielding equal or greater numbers of correct matches than observed (one-sided). Mann–Whitney U tests were two-sided.

#### Profiling drug-induced protein spatial responses

For perturbation profiling, we used paired experimental and *ProtiCelli* simulated MDA-MB-468 images under Paclitaxel and Vorinostat treatment, with untreated cells as controls. Of the 906 possible protein–drug pairs (453 proteins × 2 drugs), 891 had at least 20 paired cells and were retained for the primary perturbation-response analysis. Cells were segmented from a min–max-normalized composite of the DAPI, MT, and ER channels by Otsu thresholding, followed by morphological closing and hole filling; nuclear masks were obtained by Otsu thresholding DAPI within each cell mask. Forty morphological features were computed per cell, comprising 38 protein-channel features and two reference-derived covariates (cell area and nuclear-area fraction). The 38 protein-channel features spanned protein abundance and intensity-distribution statistics; nuclear–cytoplasmic partitioning; radial protein distribution across three regions defined from the normalized Euclidean distance transform of the cell mask; spatial autocorrelation (Moran’s *I* at lags 1, 2, 4, and 8); Manders and Pearson colocalization with DAPI, MT, and ER; displacement of protein intensity relative to cellular reference positions; and protein-positive object statistics. The two reference-derived covariates were excluded from model-performance evaluation because they are identical between experimental and simulated images by construction. To account for redundancy, features were clustered by the Spearman correlation of their experimentally measured perturbation effects (average linkage, merged at *ρ* 0.70), and each cluster was summarized by one representative; effective dimensionality was estimated from the participation ratio of the eigenvalue spectrum of the feature-effect correlation matrix (Table S4).

For each protein, drug, and feature, the perturbation effect was quantified independently in experimental and simulated images using Cliff’s *δ* between treated and untreated cells. Cliff’s *δ* was used rather than a ratio of medians because several profiling features take zero or negative values and because, as a rank-based statistic, it is invariant to strictly monotonic transformations and therefore less sensitive to global intensity-scale differences between experimental and simulated images.

Experimental effects were treated as ground truth, and experimental–simulated concordance was assessed by Spearman correlation. Statistical evidence for individual perturbation effects was assessed by two-sided Mann–Whitney *U* tests with Benjamini–Hochberg correction. Sampling repeatability was estimated by randomly dividing experimental cells into two halves, recomputing perturbation effects independently in each, and applying the Spearman–Brown correction to the between-half Spearman correlation. Recovery of strong perturbation responses was evaluated separately for Paclitaxel and Vorinostat: within each drug, proteins were ranked by the magnitude of the simulated effect, *|δ*_sim_|, and positives were defined as the 10% of proteins with the largest experimental effect magnitudes, *δ*_exp_ , fixing the positive prevalence and chance average precision at 0.10. Recovery was quantified by average precision and by precision and recall at fixed ranking depths, and directional agreement was evaluated among contrasts significant in both groups against a marginal-preserving null equivalent to a one-sided Fisher test on the direction contingency table. To determine how performance depended on response strength, proteins were grouped into deciles by experimental effect magnitude, and experimental– simulated Spearman correlation and directional agreement were computed within each decile. To distinguish protein-specific recovery from a generic response shared across proteins within a drug, effects were arranged as a protein × drug matrix for each feature, and the mean response within each drug was removed separately from the experimental and simulated effects; concordance was then recomputed across the residual protein effects. Significance was assessed against a null distribution generated by independently permuting protein identities within each drug 1,000 times. Experimental and simulated effects were also compared within Paclitaxel and Vorinostat separately by Spearman correlation across proteins.

#### Cell cycle stage assignment and pseudotime calculation

Cell cycle stages were determined using a two-step approach combining polar coordinate pseudotime modeling and Gaussian Mixture Model (GMM) classification.

To calculate the pseudotime, we first calculated nuclear mean intensities for GMNN (S/G2 marker) and CDT1 (G1 marker) and performed a log_10_ transformation. To account for the circular pattern of their expression throughout the cell cycle, the data were centered by fitting a circle using least-squares optimization to minimize variance in radial distances from the fitted center. Centered Cartesian coordinates were then converted to polar coordinates (*ρ, ϕ*), and cells were sorted by angular position *ϕ*. The angular coordinate was re-zeroed to align with the phase of minimum cell density (corresponding to mitosis, where cells are difficult to segment), determined by binning *ϕ* into 1000 bins and identifying the minimum. The pseudotime was calculated by normalizing and reversing the reordered angles to account for counter-clockwise progression:

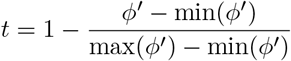

where *ϕ*^′^ represents re-zeroed angular coordinates. Finally, the pseudotime was transformed to achieve uniform cell density across the cell cycle using equal N histogram binning.

Discrete cell cycle phases were assigned using a 3-component Gaussian Mixture Model (GMM) fitted to the two-dimensional log-transformed intensity space (log_10_ GMNN, log_10_ CDT1). The GMM models the data as a mixture of three Gaussian distributions:

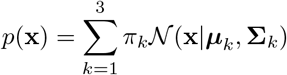

where *π_k_* are mixing coefficients, and ***µ****_k_* and **Σ***_k_* are the mean vectors and covariance matrices for each component, respectively. The three components were mapped to cell cycle phases based on their positions in the cell cycle span: G1 phase (the component with minimum mean GMNN intensity), G2 phase (the component with minimum mean CDT1 intensity), and G1/S phase (the remaining intermediate component). Each cell was assigned to the phase corresponding to the component with maximum posterior probability *p*(*k* **x***_i_*). To ensure fair comparison, images used for fine-tuning were excluded from all evaluations.

#### Pixel-level spatial clustering

For each multi-channel protein image stack, individual channels were first smoothed with a Gaussian filter (*σ* = 0.5) and binarized at 1.25× the Otsu threshold to identify protein-positive pixels. The binarized stack was then reshaped by treating channels as features and individual pixels as observations. For instance, an image stack of dimensions 100 512 512 (channels height width) was transformed into a 262, 144 100 data matrix, where each row corresponds to a single pixel’s binarized expression profile across all 100 proteins. K-means clustering (*N* = 12 clusters for subcellular partitioning examples and *N* = 8 for pathway gene set partitioning) was applied to this matrix to assign each pixel to a spatial cluster based on its characteristic protein co-localization pattern. To ensure consistent cluster identities across cells, clustering was performed globally on the combined data matrix from all images within each cell line. For 3D subcellular partitioning, each z-slice was extracted independently from 3D image stacks and combined into a single data matrix spanning all slices and all cells. Pixel-level spatial clustering was then performed on this combined matrix. Clusters were ranked by size in descending order.

#### Cluster annotation and gene set disentanglement

To assess whether spatially resolved protein occupancy reflects structured biological pathway organization, we mapped cluster expression profiles onto the Reactome hierarchy, exemplified here for the “Metabolism of lipids” pathway (R-HSA-556833) in Hep-G2 cells [61]. For each spatial cluster *k*, marker proteins were identified using a spatial enrichment score measuring the enrichment of each protein relative to all other clusters:

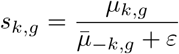

where *µ_k,g_* is the mean binarized occupancy of protein *g* in cluster *k*, *̄µ*_*k,g*_ is the unweighted mean of the mean binarized occupancies of protein *g* across the remaining clusters, and *ε* = 10^−6^ is a small constant to avoid division by zero. The background cluster was excluded from this analysis. *s_k,g_* is thus a one-vs-rest ratio between a single cluster and all other clusters, and not a ratio to the whole-cell mean. Both the absolute occupancy matrix (*µ_k,g_*) and the spatial enrichment matrix (*s_k,g_*) were computed across all proteins and clusters, and exported for downstream analysis.

#### Organelle annotation

The clusters were assigned organelle identities based on the enrichment patterns of curated marker gene sets spanning diverse organelles and subcellular structures (Table S6). For each cluster and marker set, we computed *s_k,g_* across all marker proteins of that set. Primary annotations were assigned when multiple genes within a marker set consistently exceeded *s_k,g_ >* 1.5, indicating strong specific enrichment relative to all other clusters. When no marker set reached this threshold but multiple genes from the same organelle showed consistent directional enrichment, secondary annotations were assigned based on the combined enrichment score (*µ_k,g_ × s_k,g_*) of top-ranked genes; these reflect weaker or partial spatial associations. Clusters exhibiting co-enrichment across multiple marker sets received composite identities (e.g., “ER/Golgi”), as organelles overlap in 2D projections. Clusters without consistent enrichment remained unassigned.

#### Gene set mapping

The resulting expression matrices were then mapped onto the Reactome pathway hierarchy. The Reactome ContentService REST API (https://reactome.org/ ContentService) was queried to retrieve all sub-pathways of a selected root pathway (R-HSA-556833) down to a hierarchical depth of two levels via traversal of the hasEvent relationship. Gene members of each sub-pathway were obtained via the /data/participants/ stId endpoint and extracted from ReferenceGeneProduct entries within the returned refEntities records.

For each (sub-pathway *P* , spatial cluster *k*) pair, we computed the pathway-level spatial enrichment

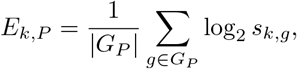

where *G_P_* is the set of proteins annotated to *P* that are present in the occupancy matrix, and *s_k,g_* is the protein-level spatial enrichment score defined above. Values of log_2_ *s_k,g_* were truncated below at 2 before averaging and before testing. Statistical significance was assessed using a one-sided Wilcoxon signed-rank test applied to the member values log_2_ *s_k,g_* : *g G_P_* (*H*_1_: median *>* 0, i.e., sub-pathway enriched in cluster *k* relative to all the clusters); pairs with fewer than four detected member proteins were assigned *p* = 1. Raw p-values were corrected for multiple testing across all (sub-pathway × cluster) pairs jointly using the Benjamini–Hochberg procedure. Sub-pathway–cluster pairs were retained for visualization if the BH-corrected p-value was below 0.05.

## Data availability

The Human Protein Atlas images used for model training and evaluation were obtained from the Subcellular section of the Human Protein Atlas, version 24, and are publicly available through the Human Protein Atlas. The CM4AI immunofluorescence images of MDA-MB-468 cells were obtained from the CM4AI dataset, Beta release, June 2025, available at https://dataverse.lib.virginia.edu/dataset.xhtml?persistentId=doi:10.18130/V3/F3TD5R.

The newly generated *FUCCI-CellCycle* dataset, including images of A-549 and MDA-MB-468 cells, and the three-dimensional microscopy images used for subcellular segmentation will be deposited in bioimage archive before publication. The *Proteome2Cell* dataset, comprising ProtiCelli-generated images for 12,800 protein entries across 2,400 cells from 12 human cell lines, will also be deposited in bioimage archive before publication and part of it will be accessible through the Human Protein Atlas virtual-cell interface upon release of Human Protein Atlas version 26.

## Code availability

All analyses were implemented in Python 3.10. Numerical operations: NumPy 1.24, SciPy 1.11. Deep learning: PyTorch 2.6.0, torchvision 0.21.0, diffusers 0.36.0. Machine learning and statistics: scikit-learn 1.8.0. Image processing: scikit-image 0.26.0. Visualization and dimensionality reduction: scanPy 1.10.2, umap-learn 0.5.

Code is available at https://github.com/CellProfiling/ProtiCelli.

## Author contributions

H.S., W.O., and E.L. conceptualized the project and designed the study. H.S. and M.L. trained machine learning models. H.S., K.K., J.N.H., W.L., W.F., and E.L. analyzed and interpreted data. H.S., K.K., and F.B. worked on code release and documentation. H.S., K.K., and J.N.H. developed tool applications, visualized results, and prepared figures. H.S., F.B., U.A. and E.L. worked on developing the HPA virtual cell portal. E.L. supervised the project. All authors edited and commented on the manuscript.

## Supplementary information

Please see our supplementary figures 1 - 11, supplementary tables 1 - 7, and supplementary video 1 in separate files.

