## Supplementary figures and tables for "Generative machine learning unlocks the first proteome-wide image of human cells"

**Supplementary Information**

Huangqingbo Sun, Konstantin Kahnert, Jan N. Hansen, William Leineweber, Mingyang Li, Wanyue Feng, Frederic Ballllosera Navarro, Ulrika Axelsson, Wei Ouyang, Emma Lundberg

September 1, 2026

### List of Supplementary Tables

**Table S1.** Protein and cell line names supported by *ProtiCelli. External spreadsheet.*

**Table S2.** Drug-perturbation response statistics for *ProtiCelli*-generated and real immunofluorescence images. *External spreadsheet.*

**Table S3.** Per-protein drug-perturbation effects for the radial intensity feature (*ProtiCelli*-generated versus real images). *External spreadsheet.*

**Table S4.** Feature-level recovery summary across the 40 morphological features.

**Table S5.** FUCCI marker intensity statistics across cell-cycle stages.

**Table S6.** Curated organelle marker proteins for subcellular compartment segmentation.

**Table S7.** Spatial enrichment of lipid-metabolism proteins across *ProtiCelli*-defined clusters in Hep-G2 cells. *External spreadsheet.*

### 1 Supplementary Figures

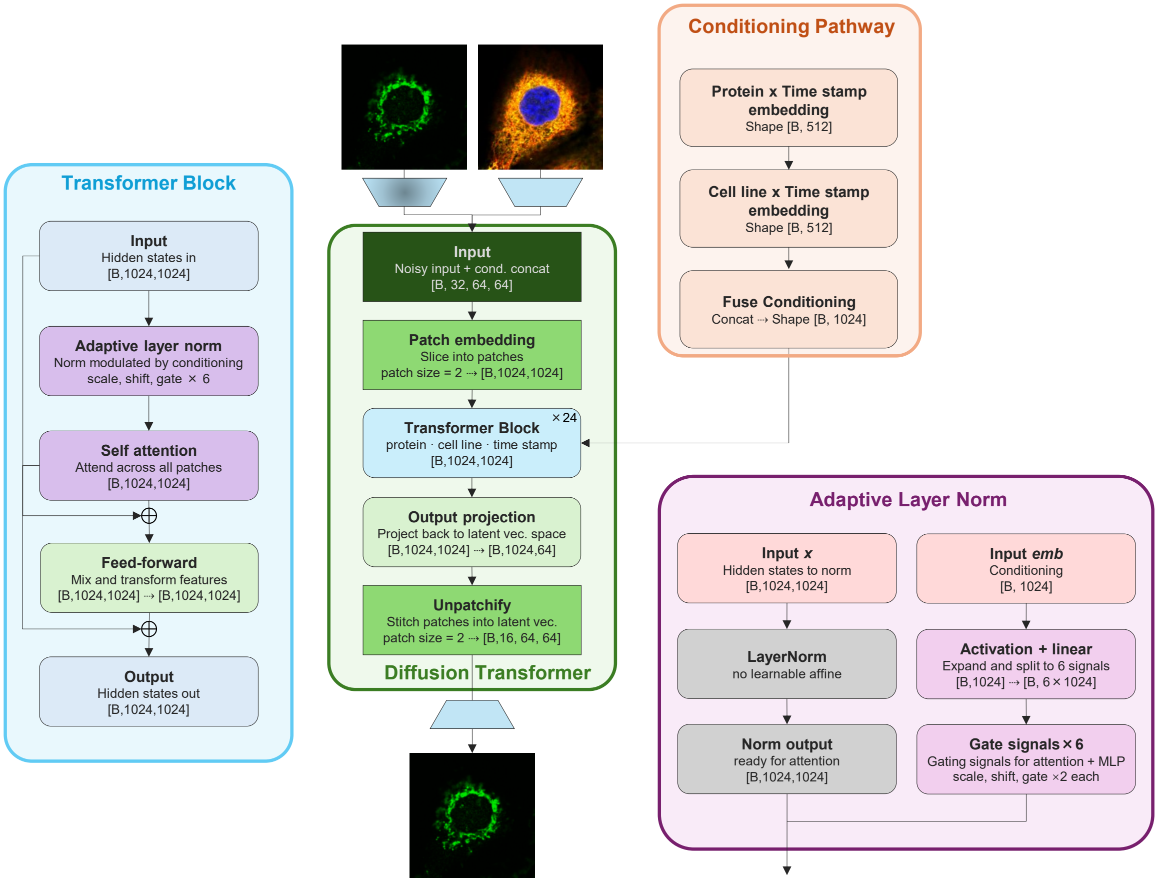

Figure S1: **Neural network architecture of *ProtiCelli*, a diffusion transformer for conditional cell image generation.** The model (middle) takes a noisy latent as input and denoises it through a stack of 24 transformer blocks, conditioned jointly on protein identity, cell line identity, and diffusion timestep. The conditioning embeddings are fused and injected into every block, and the output is projected and reassembled into the final denoising prediction in latent space. Each transformer block (left) applies self-attention and a feed-forward network in sequence, each wrapped in residual connections; conditioning is injected via an adaptive layer norm that predicts scale, shift, and gate signals from the joint conditioning embedding (right) to modulate both the attention and feed-forward sub-layers.

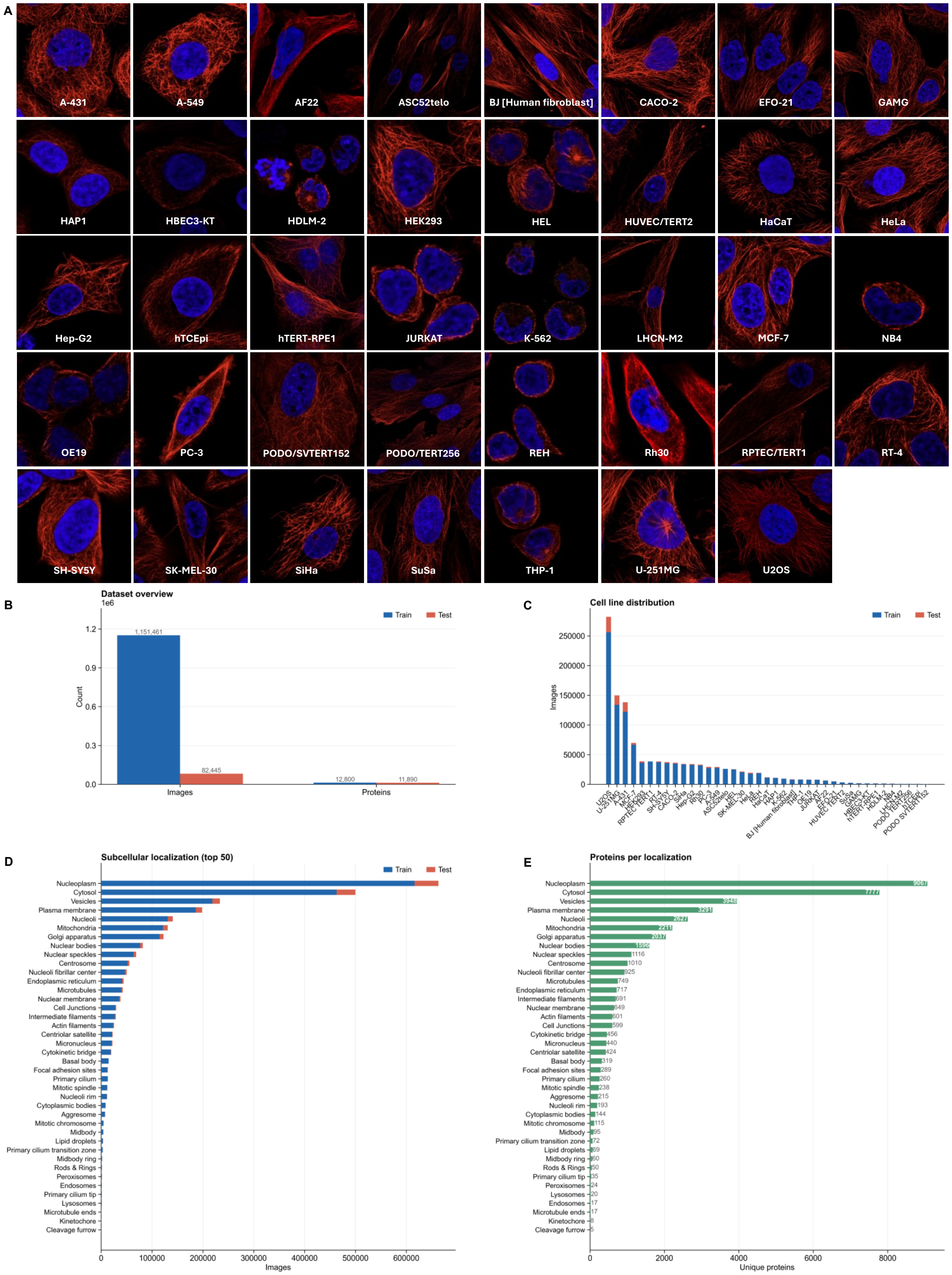

Figure S2: **Representative immunofluorescence images of the 39 cell lines and overview of the *ProtiCelli* training and test datasets.** (Caption continued on next page.)

Figure S2: **Representative immunofluorescence images of the 39 cell lines (continued).** (**A**) Each panel shows a single cell stained for microtubules (red) and nuclei (DAPI, blue), with cell line identity labeled below. The collection spans diverse morphologies, tissue origins (>20 tissues), and biological contexts, encompassing cancerous and non-cancerous lines of epithelial, mesenchymal, myeloid, and endothelial origin. (**B**) Total number of single-cell images (1,151,461 train; 82,445 test) and unique proteins (12,800 train; 11,890 test). (**C**) Per-cell-line image counts for train and test splits across all 39 cell lines, sorted by total image count; the U2OS line is the most represented. (**D**) Image counts stratified by subcellular localization annotation for the top 50 compartments; nucleoplasm, cytosol, and plasma membrane are the most abundant. (**E**) Number of unique proteins per subcellular localization compartment; nucleoplasm contains the largest number of distinct proteins (9,069), followed by cytosol (7,781), and vesicles (3,949).

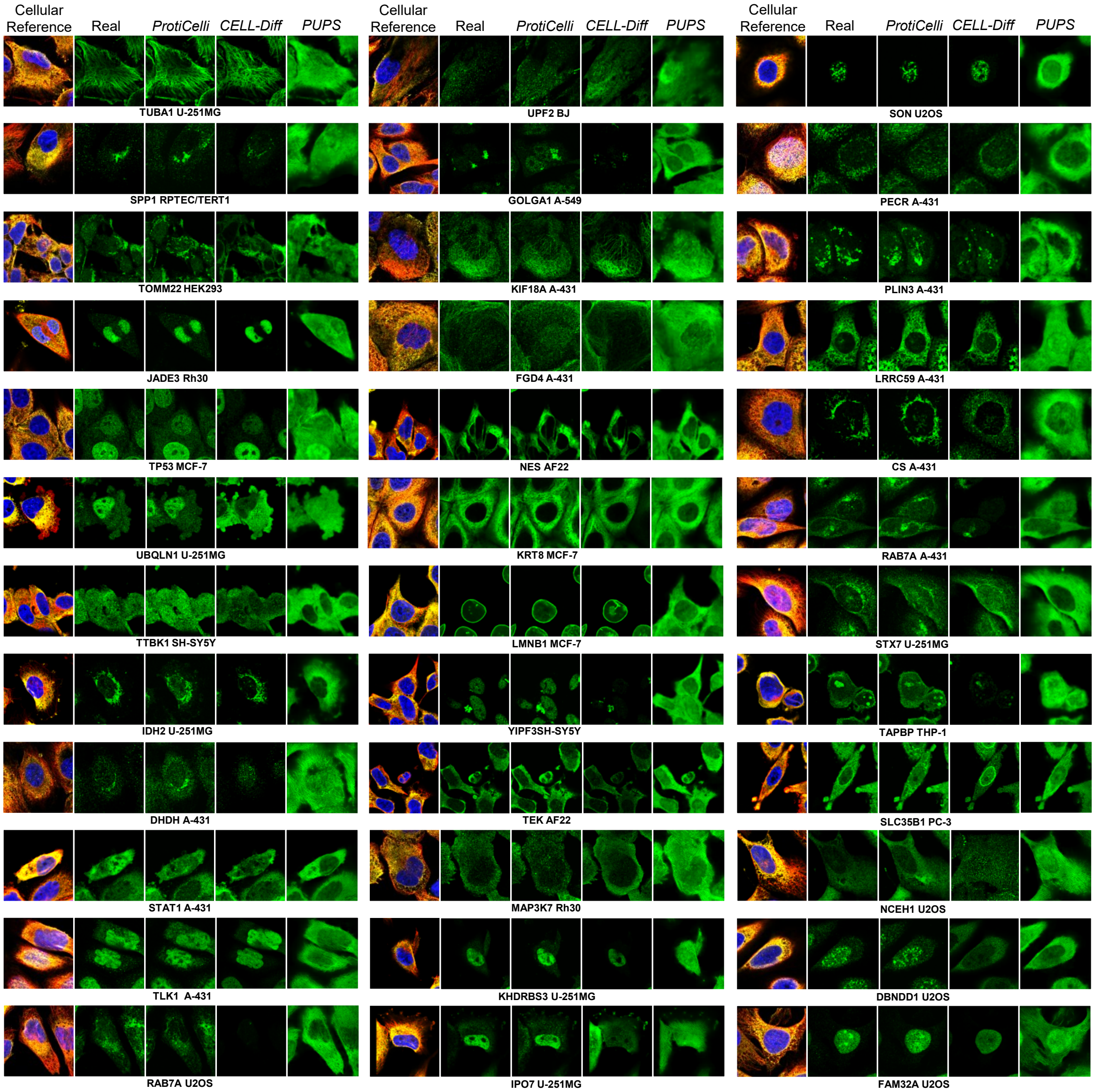

Figure S3: **Qualitative comparison of fluorescence microscopy images generated by *ProtiCelli*, *CELL-Diff*, and *PUPS* across diverse proteins and cell lines.** Each row shows one protein in a given cell line (labeled below). Columns, left to right: the cellular reference (microtubules, red; nucleus, blue), the real protein channel (green), and predictions from *ProtiCelli*, *CELL-Diff*, and *PUPS*. Examples span a range of subcellular localizations, morphologies, and cell lines.

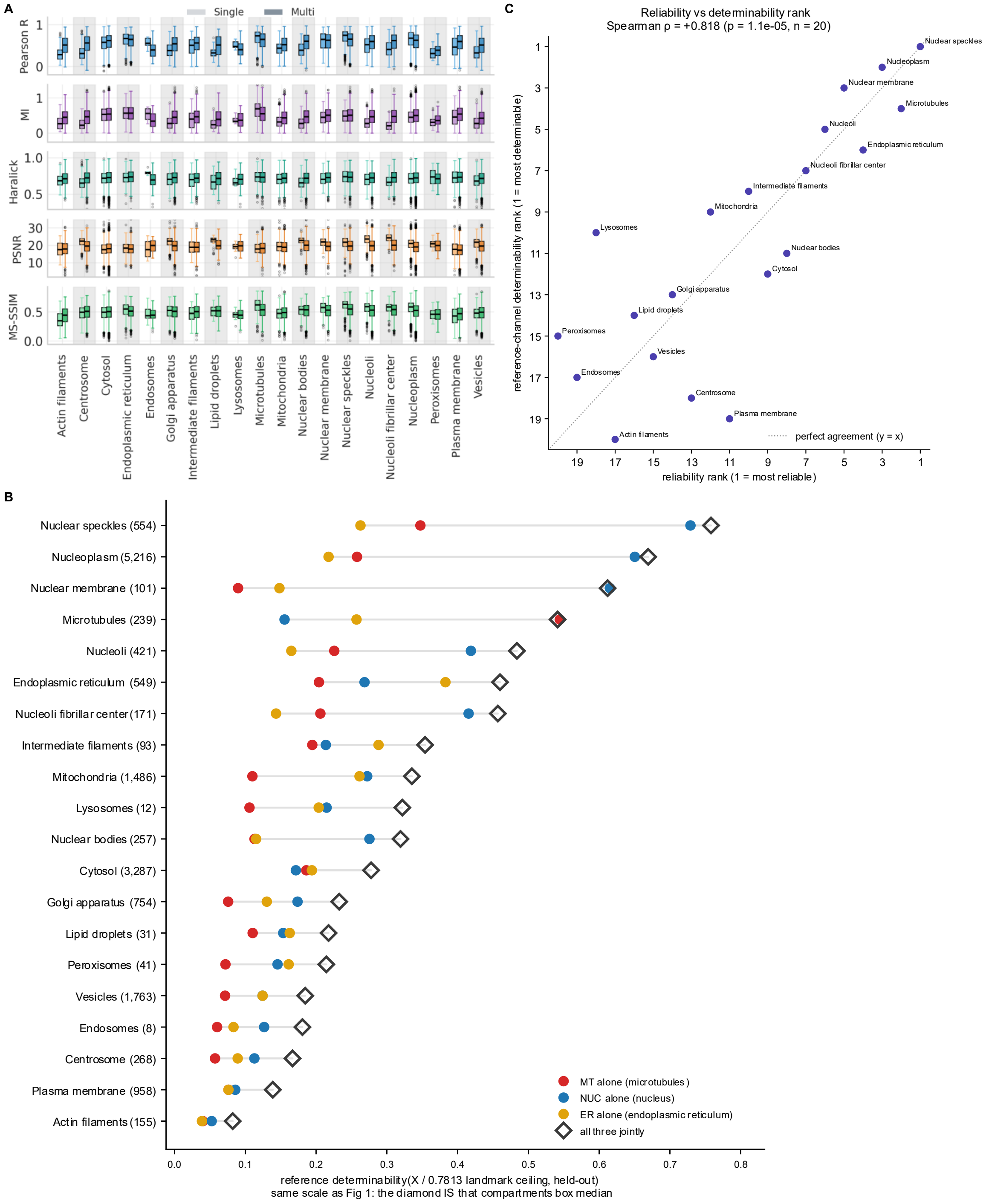

Figure S4: Captions on the next page.

Figure S4: **Evaluation of *ProtiCelli* performance for single- versus multi-localizing proteins, and rank correlation between organelle reliability and determinability.** (**A**) Per-organelle prediction performance for single-localized (light) versus multi-localized (dark) proteins, scored by five image-similarity metrics (Pearson $r$, mutual information, Haralick texture correlation, PSNR, MS-SSIM; rows) across 20 subcellular compartments (columns). Multi-localized proteins score comparably to or above single-localized proteins in most compartments, plausibly because at least one predominant localization falls in a well-represented major organelle, and strong prediction there compensates for weaker performance at secondary, less common sites. (**B**) Determinability for each subcellular compartment, computed from one reference stain alone (circles: MT, microtubules; NUC, nucleus; ER, endoplasmic reticulum) or from all three together (open diamonds). X is the fraction of a protein's within-cell positional information recoverable from the reference stains, normalized so that 1 is the maximum this method can resolve (Methods). Points are compartment medians across single-localizing proteins; n = protein–cell-line pairs per compartment, shown in parentheses (16,364 pairs, 8,474 proteins, 12 cell lines). Values are on the same scale as Fig. 2I, in which the diamond is the compartment median. (**C**) Per-organelle determinability rank versus mean *ProtiCelli* reliability rank across 20 subcellular compartments. Ranks are strongly correlated (Spearman $\rho = 0.827$, $p = 6.9 \times 10^{-6}$, $n = 20$); the dotted line indicates perfect rank agreement ($y = x$).

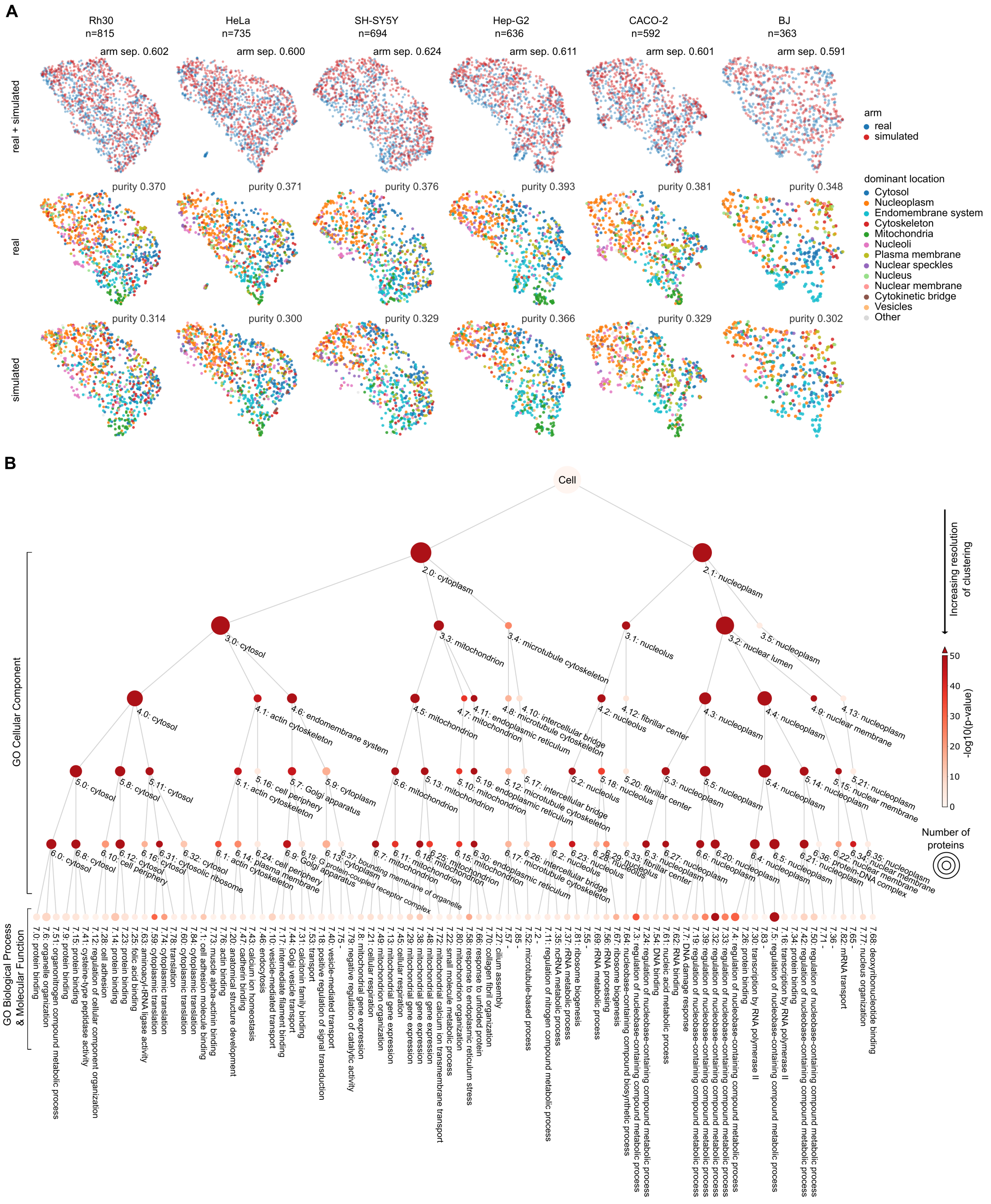

Figure S5: **Quantitative and qualitative evaluation of *ProtiCelli* SubCell embeddings.** (**A**) UMAP projections of *SubCell* embeddings for the six HPA cell lines with the least stained proteins; the remaining six are shown in Fig. 3A. Each point represents one protein. Real and simulated embeddings were mean-centered and jointly projected for each cell line (Methods). Columns denote cell lines; the top row shows embeddings colored by image source, and the lower rows show real and simulated embeddings separately in the same projection, colored by primary subcellular location. Separation and purity were calculated using the full 1,536-dimensional embeddings. **(B)** Multiscale hierarchical map obtained by iterative subclustering of 9,543 protein representations derived from real HPA images. Clusters and subclusters are annotated using enriched Gene Ontology Cellular Component terms at levels 1–6 and Gene Ontology Biological Process or Molecular Function terms at level 7; the most significantly enriched term is shown for each node. Node size indicates the number of proteins, and node color indicates the $-\log_{10}$-transformed enrichment $p$-value. Figure reproduced from [43].

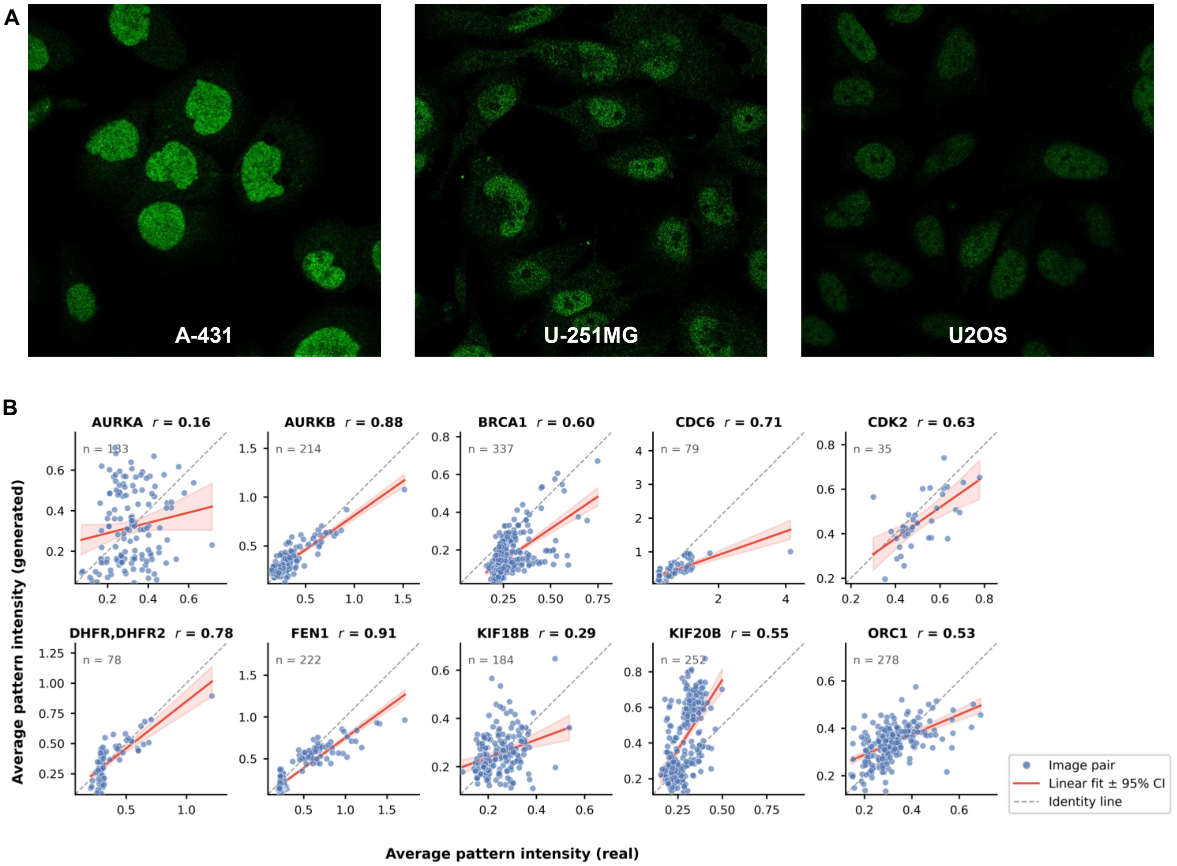

Figure S6: ***ProtiCelli* recovers cell-cycle-dependent expression patterns.** (**A**) Representative real immunofluorescence images of BPTF from the Human Protein Atlas across three cell lines (A-431, U-251MG, U2OS). (**B**) Scatter plots comparing the average pattern intensity of *ProtiCelli*-generated images against matched real fluorescence images for ten further cell-cycle regulated proteins. Each point is one image pair; $n$ pairs per protein are indicated. The red line denotes the linear fit with 95% confidence interval; the dashed gray line indicates the identity ($y = x$). Pearson correlation coefficients ($r$) are reported per protein.

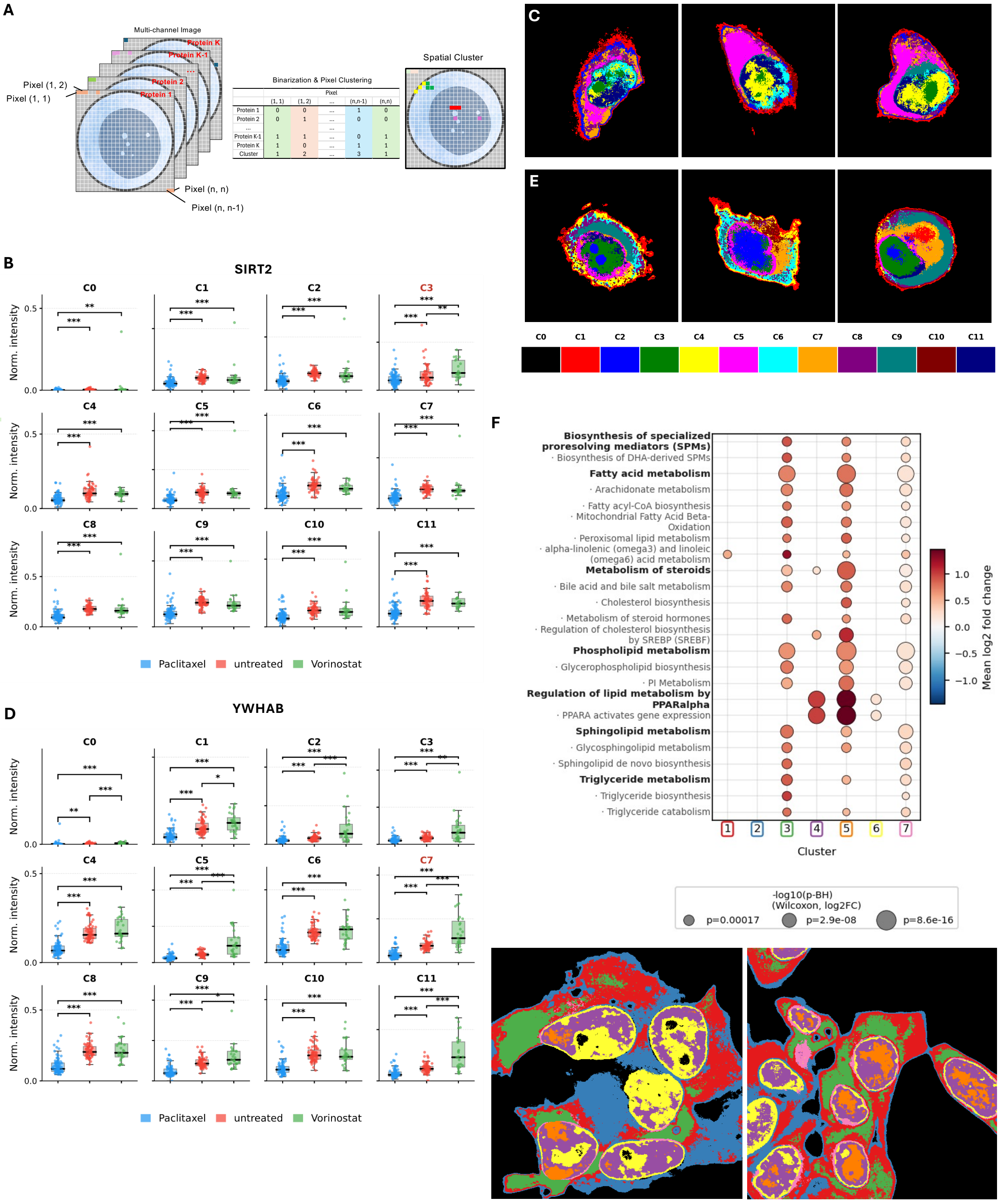

Figure S7: Captions on the next page.

Figure S7: **Unsupervised spatial clustering, subcellular region partition, and intensity quantification and spatial localization** (**A**) (*Left*) A multi-channel fluorescence microscopy image is represented as a stack of $K$ single-protein intensity maps, each encoding the spatial expression pattern of a distinct protein across an $n \times n$ pixel grid. (*Center*) Each channel is independently binarized per pixel (1 = expressed, 0 = absent), producing a binary co-expression vector of length $K$ per pixel; pixels are then grouped into discrete clusters based on the similarity of these vectors. (*Right*) The resulting spatial cluster map color-codes each pixel by its assigned cluster, revealing compartments defined by shared protein co-expression signatures. This unsupervised approach derives spatial compartmentalization directly from *ProtiCelli* generated images, without relying on predefined protein groups or anatomical annotations. (**B**) Normalized SIRT2 intensity across 12 unsupervised spatial clusters (C0–C11) under three treatment conditions: Paclitaxel, Vorinostat, and untreated. Boxplots display the distribution of mean intensities, normalized globally across all conditions. Statistical significance between conditions is indicated by bracket annotations ($^{*}p < 0.05$, $^{**}p < 0.01$, $^{***}p < 0.001$). Cluster 3 (red label) identifies the predicted nucleoli compartment, which shows the highest SIRT2 enrichment. (**C**) Representative spatial cluster maps for three individual MDA-MB-468 cells, color-coded according to the cluster legend (C0–C11, bottom). (**D**) Equivalent analysis for YWHAB intensity across the same 12 clusters; Cluster 7 (red label) denotes the primary YWHAB-enriched compartment. (**E**) Representative spatial cluster maps for three individual cells for YWHAB, as in (**C**). In all panels, each pixel is assigned to its corresponding co-expression cluster derived from *ProtiCelli*-generated images. (**F**) As in Figure 5G, but with both spatial clustering and enrichment analysis performed exclusively using *ProtiCelli*-generated pathway protein images, rather than organelle marker images. Spatial cluster maps of two representative cells are shown below the dot plot (7 clusters; color-coded by cluster identity).

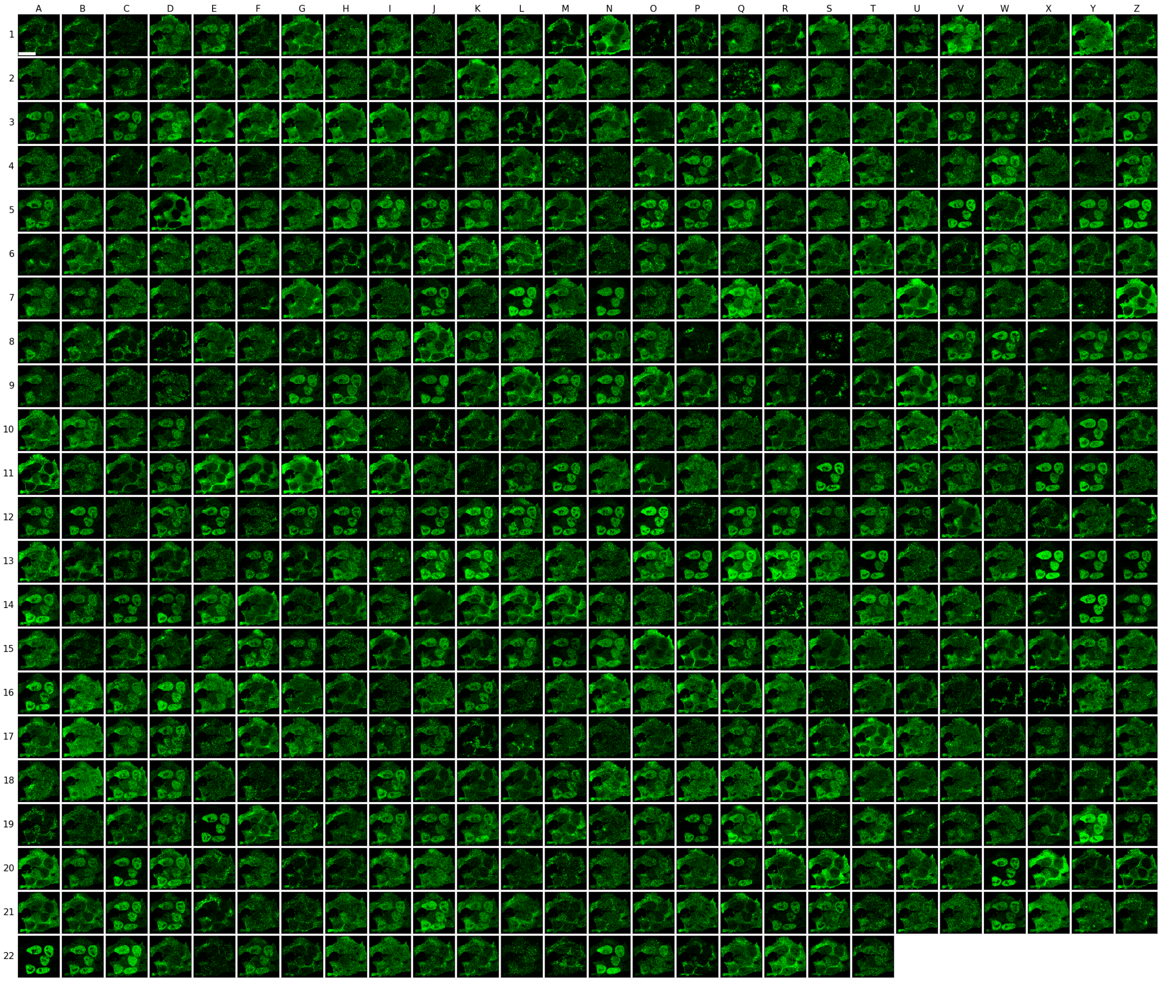

Figure S8: ***ProtiCelli* generates nearly complete lipid-metabolism proteome in a single Hep-G2 cell.** Predicted fluorescence images for all 566 proteins in the lipid metabolism pathway, generated on one fixed Hep-G2 reference cell. Tiles are arranged row-major (columns A–Z, rows 1–22) in the order listed in Table S7; trailing positions are blank. Pixel-level spatial clustering of these predictions partitions the cell into distinct compartments enriched for unique biological functions (Fig. 5G). Scale bar, 20 $\mu m$.

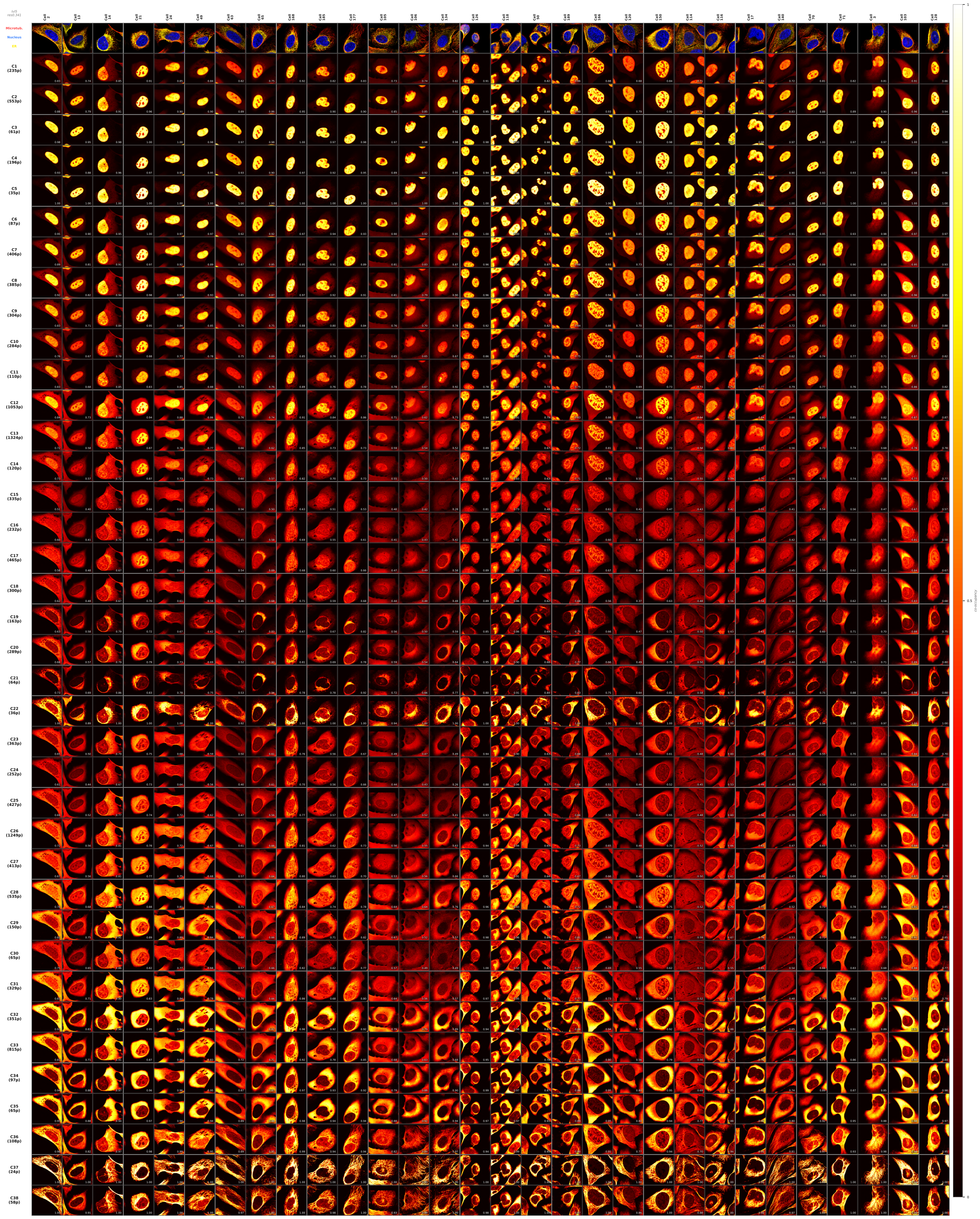

Figure S9: **Cluster co-occupancy patterns are conserved across cells.** Consensus expression heatmaps for each cluster of the hierarchical multi-scale map (Fig. 6B), computed from *ProtiCelli*-generated images and compared across cells. Columns are individual cells; the reference cell of Fig. 6B is Cell 134 here. The top row shows each cell's reference channel (microtubules, red; nucleus, blue). Remaining rows are clusters C1–C38 (top to bottom), with per-cluster protein count in parentheses; cluster IDs match Fig. 6B, and resolution increases down the hierarchy. Heatmap images visualize the consensus expression pattern of all proteins in each cluster, derived from *ProtiCelli*-generated images of the same cell (brighter color indicates higher overlap of protein expression, see color bar on the right, which indicates the Co-occupancy fraction). The number annotated in each panel indicates the maximal intensity of the heatmap.

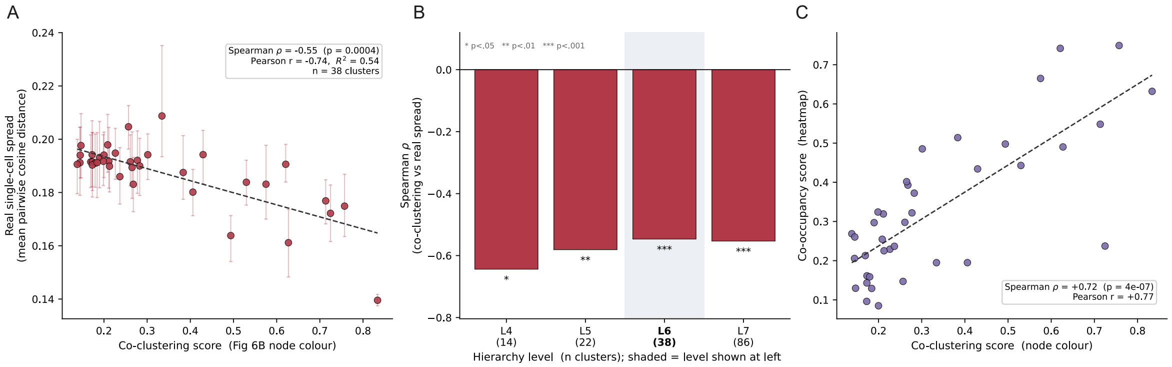

Figure S10: **Model-derived cluster stability scores predict single-cell spatial cohesion in real images.** **(A)** For each of the 38 leaf clusters, higher model co-clustering scores correlate with tighter real protein spread, measured as the mean pairwise cosine distance of the cluster's proteins in the real-image SubCell embedding space (Spearman $\rho = -0.55$, $p = 4 \times 10^{-4}$; Pearson $r = -0.74$). Spread is computed at a fixed protein budget to control cluster size; error bars are the 95% bootstrap confidence interval (2.5th–97.5th percentile over 100 resamples). **(B)** The negative correlation is robust across depths of the clustering hierarchy (14–86 clusters); stars denote per-level Spearman significance ($^*p < 0.05$, $^{**}p < 0.01$, $^{***}p < 0.001$). **(C)** The model's two predictive metrics—co-clustering (pattern similarity across individual cells) and co-occupancy (spatial protein overlap within each cluster)—strongly agree ($\rho = +0.72$). Generated embeddings were used only to define clusters, never to measure spread.

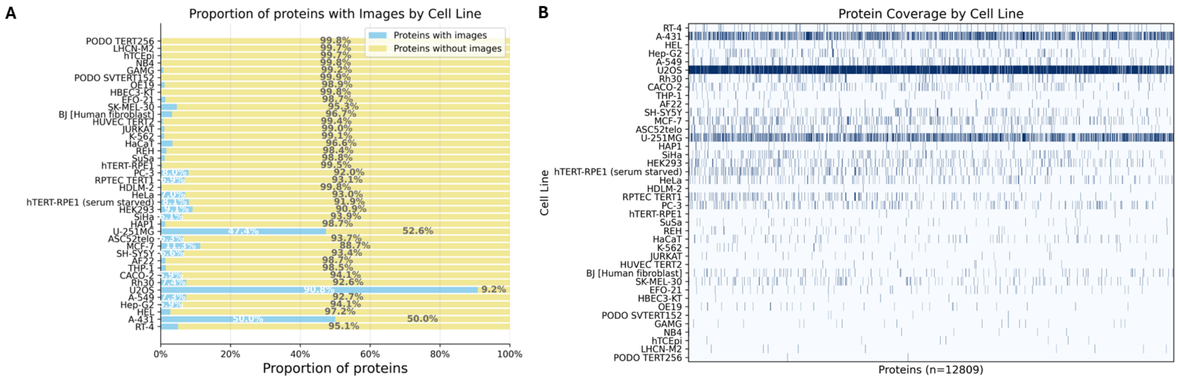

Figure S11: **Sparsity within the HPA subcellular image dataset.** (**A**) Distribution of protein coverage across cell lines. Only three cell lines contain images for approximately more than half of the proteins. (**B**) Distribution of cell line coverage across proteins. Most proteins are imaged in no more than three cell lines, demonstrating substantial sparsity.

#### 2 Supplementary Tables

Table S1. **Protein and cell line names supported by *ProtiCelli*.** Each row lists one of the 12,800 unique proteins included in the training dataset alongside the cell line in which it was imaged. The dataset spans 39 cell lines: A-431, A-549, AF22, ASC52telo, BJ [Human fibroblast], CACO-2, EFO-21, GAMG, HAP1, HBEC3-KT, HDLM-2, HEK293, HEL, HUVEC/TERT2, HaCaT, HeLa, Hep-G2, hT-CEpi, hTERT-RPE1, JURKAT, K-562, LHCN-M2, MCF-7, NB4, OE19, PC-3, PODO/SVTERT152, PODO/TERT256, REH, Rh30, RPTEC/TERT1, RT-4, SH-SY5Y, SiHa, SK-MEL-30, SuSa, THP-1, U-251MG, and U2OS.

Table S2. **Drug-perturbation response statistics for *ProtiCelli*-generated and real immunofluorescence images.** Each row corresponds to one of 453 proteins. For each drug condition (Paclitaxel, Vorinostat), the following quantities are reported for both real and *ProtiCelli*-generated images: statistical significance of the intensity difference between treated and untreated cells (Sig; ns, *, **, ***), direction of change (Dir; up or down), and the associated p-value (Mann-Whitney U test). The final column per drug (DirMatch; Y/N) indicates whether the direction of change is concordant between real and generated images.

Table S3. **Per-protein drug-perturbation effects for the radial intensity feature (*ProtiCelli*-generated versus real images).** Each row is one of 453 proteins under one drug condition (Paclitaxel or Vorinostat), for the 891 protein $\times$ condition contrasts with at least 20 paired cells. The effect is Cliff's $\delta$ of the innermost-shell radial intensity fraction (`rad0`) between treated and untreated cells, computed independently on real ($d_{\text{real}}$) and *ProtiCelli*-generated ($d_{\text{pred}}$) images; positive values denote redistribution of signal toward the cell centre. $q_{\text{real}}$ and $q_{\text{pred}}$ are Benjamini-Hochberg-adjusted Mann-Whitney $p$-values computed within each protein. DirMatch (Y/N) indicates whether the sign of Cliff's $\delta$ agrees between the two arms; $n_{\text{cells}}$ is the number of paired treated cells. This table reports the continuous, scale-invariant effect measure underlying Figure 4; it is distinct from the binarized-pattern significance analysis of Table S2 and the two are not expected to agree protein-by-protein.

Table S4: **Feature-level recovery summary across the 40 morphological features.** Ranked by attenuation-corrected concordance (Frac), the real-versus-*ProtiCelli* Spearman correlation as a fraction of the split-half repeatability ceiling of the experimental measurement. $\rho$, raw real-versus-generated Spearman across all contrasts; Ceiling, split-half repeatability; Dir and Fold, direction agreement and its ratio to a marginal-preserving hypergeometric null; $p_{\text{dir}}$, that null's $p$-value. Flag marks features that are not scale-invariant (ABS; biased by the decoder intensity clamp, excluded from primary evaluation) or reference-derived and identical between arms by construction (COVAR; covariates, never evaluation targets). Cluster gives the representative of each redundancy cluster ($|\rho| \geq 0.70$ on measured effects); features sharing a representative are not independent. The radial features form a single cluster (effective dimensionality 4.2 of 40).

| Feature | Flag | Cluster | $\rho$ | Ceiling | Frac | Dir | Fold | $p_{\text{dir}}$ |
|---|---|---|---|---|---|---|---|---|
| area_cell | COVAR | area_cell | 1.00 | 0.86 | 1.16 | 1.00 | 1.16 | $3e-58$ |
| nuc_area_frac | COVAR | nuc_area_frac | 1.00 | 0.89 | 1.13 | 1.00 | 1.26 | $2e-90$ |
| rad2 | – | rad0 | 0.62 | 0.79 | 0.78 | 0.95 | 1.25 | $3e-31$ |
| rad_com | – | rad0 | 0.59 | 0.84 | 0.71 | 0.91 | 1.17 | $3e-23$ |
| rad0 | – | rad0 | 0.56 | 0.84 | 0.66 | 0.91 | 1.19 | $4e-22$ |
| nuc_frac | – | nuc_frac | 0.59 | 0.89 | 0.66 | 0.93 | 1.16 | $6e-28$ |
| rad1 | – | rad0 | 0.55 | 0.84 | 0.66 | 0.90 | 1.22 | $6e-20$ |
| total_cell | ABS | total_cell | 0.55 | 0.93 | 0.59 | 0.77 | 1.59 | $1e-32$ |
| mass_disp | – | mass_disp | 0.41 | 0.71 | 0.57 | 0.93 | 1.71 | $2e-19$ |
| n_obj | – | n_obj | 0.45 | 0.87 | 0.52 | 0.95 | 1.01 | $4e{-3}$ |
| manders_atub | – | manders_atub | 0.46 | 0.88 | 0.52 | 0.81 | 1.52 | $2e-23$ |
| manders_dapi | – | nuc_frac | 0.43 | 0.88 | 0.50 | 0.87 | 1.19 | $2e-19$ |
| mean_all | ABS | total_cell | 0.46 | 0.93 | 0.49 | 0.74 | 1.51 | $4e-24$ |
| manders_er | – | manders_atub | 0.41 | 0.90 | 0.45 | 0.79 | 1.34 | $5e-18$ |
| total | ABS | total_cell | 0.41 | 0.93 | 0.44 | 0.73 | 1.41 | $2e-19$ |
| obj_area_cv | – | obj_area_cv | 0.31 | 0.78 | 0.40 | 0.77 | 1.46 | $3e{-9}$ |
| otsu_thr | ABS | total_cell | 0.34 | 0.93 | 0.36 | 0.79 | 1.12 | $4e{-9}$ |
| corr_dapi | – | corr_atub | 0.32 | 0.91 | 0.35 | 0.71 | 1.46 | $1e-16$ |
| mean_otsu | ABS | total_cell | 0.30 | 0.92 | 0.33 | 0.79 | 1.10 | $9e{-7}$ |
| corr_er | – | corr_atub | 0.28 | 0.90 | 0.31 | 0.71 | 1.39 | $2e-11$ |
| mean_cell | ABS | total_cell | 0.29 | 0.94 | 0.31 | 0.82 | 1.05 | $1e{-4}$ |
| p99_cell | ABS | total_cell | 0.27 | 0.92 | 0.29 | 0.77 | 1.08 | $4e{-5}$ |
| nc_ratio | – | corr_atub | 0.25 | 0.88 | 0.28 | 0.73 | 1.46 | $2e-13$ |
| corr_atub | – | corr_atub | 0.25 | 0.90 | 0.28 | 0.74 | 1.30 | $1e-11$ |
| cv_cell | – | total_cell | 0.26 | 0.94 | 0.27 | 0.87 | 1.06 | $9e{-8}$ |
| gini | – | total_cell | 0.25 | 0.94 | 0.26 | 0.87 | 1.07 | $5e{-8}$ |
| top1_frac | – | total_cell | 0.24 | 0.94 | 0.26 | 0.87 | 1.05 | $2e{-6}$ |
| entropy | – | total_cell | 0.24 | 0.92 | 0.26 | 0.91 | 1.04 | $9e{-7}$ |
| top5_frac | – | total_cell | 0.24 | 0.94 | 0.25 | 0.87 | 1.05 | $5e{-6}$ |
| moran_lag8 | – | moran_lag8 | 0.22 | 0.89 | 0.25 | 0.78 | 1.05 | $8e{-3}$ |
| skew | – | total_cell | 0.23 | 0.92 | 0.25 | 0.88 | 1.04 | $2e{-5}$ |
| p90_p10 | – | total_cell | 0.21 | 0.93 | 0.23 | 0.84 | 1.07 | $2e{-6}$ |
| kurt | – | total_cell | 0.21 | 0.92 | 0.23 | 0.87 | 1.04 | $8e{-5}$ |
| p90_p50 | – | total_cell | 0.19 | 0.93 | 0.21 | 0.85 | 1.06 | $2e{-6}$ |
| moran_lag4 | – | moran_lag8 | 0.18 | 0.90 | 0.20 | 0.72 | 1.04 | $6e{-2}$ |
| mad_ratio | – | total_cell | 0.19 | 0.93 | 0.20 | 0.81 | 1.07 | $3e{-5}$ |
| obj_area | – | moran_lag8 | 0.15 | 0.87 | 0.18 | 0.76 | 1.04 | $2e{-2}$ |
| moran_lag2 | – | moran_lag8 | 0.14 | 0.89 | 0.16 | 0.64 | 1.09 | $9e{-3}$ |
| area_frac | – | area_frac | 0.10 | 0.93 | 0.10 | 0.83 | 1.03 | $2e{-2}$ |
| moran_lag1 | – | moran_lag1 | 0.05 | 0.88 | 0.06 | 0.50 | 1.11 | $5e{-2}$ |

Table S5: **FUCCI marker intensity statistics across cell cycle stages.** Summary statistics for normalized whole-cell CDT1 and GMNN intensity across cell cycle stages (G1, G1/S, G2) in A-549 and MDA-MB-468 cell lines, for *ProtiCelli* (with and without fine-tuning) and real immunofluorescence images. Columns report number of cells ($n$), mean, median, and standard deviation.

| **Cell Line** | **Condition** | **Marker** | **Stage** | $n$ | **Mean** | **Median** | **Std** |
|---|---|---|---|---|---|---|---|
| A-549 | No fine-tuning | CDT1 | G1 | 604 | 0.439 | 0.432 | 0.107 |
| A-549 | No fine-tuning | CDT1 | G1/S | 623 | 0.437 | 0.435 | 0.104 |
| A-549 | No fine-tuning | CDT1 | G2 | 574 | 0.441 | 0.440 | 0.097 |
| A-549 | No fine-tuning | GMNN | G1 | 601 | 0.308 | 0.299 | 0.118 |
| A-549 | No fine-tuning | GMNN | G1/S | 625 | 0.316 | 0.307 | 0.114 |
| A-549 | No fine-tuning | GMNN | G2 | 578 | 0.325 | 0.311 | 0.130 |
| A-549 | Fine-tuned | CDT1 | G1 | 604 | 0.226 | 0.149 | 0.184 |
| A-549 | Fine-tuned | CDT1 | G1/S | 623 | 0.197 | 0.117 | 0.183 |
| A-549 | Fine-tuned | CDT1 | G2 | 574 | 0.151 | 0.083 | 0.162 |
| A-549 | Fine-tuned | GMNN | G1 | 601 | 0.147 | 0.057 | 0.203 |
| A-549 | Fine-tuned | GMNN | G1/S | 625 | 0.146 | 0.053 | 0.212 |
| A-549 | Fine-tuned | GMNN | G2 | 578 | 0.427 | 0.518 | 0.291 |
| A-549 | Real | CDT1 | G1 | 604 | 0.305 | 0.272 | 0.143 |
| A-549 | Real | CDT1 | G1/S | 623 | 0.185 | 0.204 | 0.087 |
| A-549 | Real | CDT1 | G2 | 574 | 0.196 | 0.215 | 0.089 |
| A-549 | Real | GMNN | G1 | 601 | 0.208 | 0.238 | 0.119 |
| A-549 | Real | GMNN | G1/S | 625 | 0.196 | 0.229 | 0.120 |
| A-549 | Real | GMNN | G2 | 578 | 0.500 | 0.526 | 0.224 |
| MDA-MB-468 | No fine-tuning | CDT1 | G1 | 377 | 0.487 | 0.488 | 0.182 |
| MDA-MB-468 | No fine-tuning | CDT1 | G1/S | 77 | 0.352 | 0.353 | 0.179 |
| MDA-MB-468 | No fine-tuning | CDT1 | G2 | 462 | 0.470 | 0.462 | 0.182 |
| MDA-MB-468 | No fine-tuning | GMNN | G1 | 379 | 0.498 | 0.495 | 0.111 |
| MDA-MB-468 | No fine-tuning | GMNN | G1/S | 78 | 0.462 | 0.461 | 0.134 |
| MDA-MB-468 | No fine-tuning | GMNN | G2 | 462 | 0.521 | 0.510 | 0.116 |
| MDA-MB-468 | Fine-tuned | CDT1 | G1 | 377 | 0.455 | 0.430 | 0.112 |
| MDA-MB-468 | Fine-tuned | CDT1 | G1/S | 77 | 0.431 | 0.389 | 0.165 |
| MDA-MB-468 | Fine-tuned | CDT1 | G2 | 462 | 0.391 | 0.371 | 0.091 |
| MDA-MB-468 | Fine-tuned | GMNN | G1 | 379 | 0.111 | 0.051 | 0.161 |
| MDA-MB-468 | Fine-tuned | GMNN | G1/S | 78 | 0.269 | 0.181 | 0.230 |
| MDA-MB-468 | Fine-tuned | GMNN | G2 | 462 | 0.511 | 0.551 | 0.214 |
| MDA-MB-468 | Real | CDT1 | G1 | 377 | 0.167 | 0.140 | 0.084 |
| MDA-MB-468 | Real | CDT1 | G1/S | 77 | 0.198 | 0.124 | 0.192 |
| MDA-MB-468 | Real | CDT1 | G2 | 462 | 0.095 | 0.097 | 0.032 |
| MDA-MB-468 | Real | GMNN | G1 | 379 | 0.175 | 0.152 | 0.112 |
| MDA-MB-468 | Real | GMNN | G1/S | 78 | 0.288 | 0.304 | 0.125 |
| MDA-MB-468 | Real | GMNN | G2 | 462 | 0.530 | 0.516 | 0.145 |

Table S6: **Curated organelle marker proteins for subcellular compartment segmentation.** A collection of marker proteins associated with common organelles and subcellular compartments, selected based on established localization annotations from the Human Protein Atlas. *ProtiCelli*-generated images of these markers were used for pixel-level spatial clustering to define spatially distinct compartments.

| **Organelle** | **Proteins** |
|---|---|
| Nuclear membrane | TPR, LMNB1, SUN2, TMPO, LEMD2, TOR1AIP1, LBR, LMNB2, NUP153, CASC3 |
| Nucleoli / rim | DDX47, RPF1, UTP6, NOL10, FTSJ3, UBTF, NAA50, NOP53, NOP56, NOLC1, POLR2K, NOP16, UTP4, NOP2, RSL1D1, MKI67 |
| Nucleoplasm | RPS19, TAF15, SMARCAD1, RAN, HNRNPA1, H2AZ1, HNRNPC, HMGB1, HNRNPK, HNRNPA3, ATF4 |
| Nuclear speckles / bodies | SRRM2, RBM25, PML, CENPC, MORF4L1 |
| Actin filaments / focal adhesions | SEPTIN9, FGD4, ZYX, VCL, GSN, MTSS2, TNS3, LAD1, LIMCH1, SPECC1L, TNS1, ARSJ |
| Microtubules | TUBA1A/TUBA1B/TUBA1C |
| Mitochondria | CS, LRPPRC, SLC25A24, TIMM44, GCDH, TRAP1, MT-CYB, ATP5F1B, HSPD1, CHCHD2, SLC25A3, COX4I1, ATP5ME, ATP5MK, TOMM6, TOMM20, TOMM40, NDUFA13 |
| Intermediate filaments | KRT19, KRT4, DES, NES, KRT17, KRT13, VIM, KRT8, KRT14, NEFM, GFAP, KRT80 |
| Cytosol | ADSL, ATXN2, G3BP2, AIMP1, SERBP1, CCDC43, ATXN2L, AMPD2, RAB-GAP1 |
| Golgi apparatus | GOLGB1, GOLGA5, GALNT2, ZFPL1, GORASP2, GOLM1, GOLIM4, CD74, SPP1, RER1, LMAN2, COPE, SDF4, TMED3, NUCB2, SRGN |
| Plasma membrane / cell periphery | STX4, SLC16A1, EZR, EPB41L3, CTNNB1, ANK3, SLC41A3, AP2M1, MSN, GNB2, CD9, SLC38A2, ATP1B3, ITGA3, BASP1, HTRA1 |
| Endoplasmic reticulum | HSP90B1, CANX, KTN1, PDIA3, RRBP1, SEC61B, CALR, P4HB, CALU, COL1A2, RTN4, RPN2, BCAP31, SSR4, RPL41 |

Table S7. **Spatial enrichment of lipid metabolism proteins across *ProtiCelli*-defined clusters in Hep-G2 cells.** This table contains five sheets: (1) Metabolism of lipid genes: the full list of 566 (supported by *ProtiCelli*) out of 758 genes in the Reactome lipid metabolism pathway (R-HSA-556833), arranged consistently with Fig. S8. (2) Expression (organelle markers): for each of the 566 pathway proteins supported by *ProtiCelli*, the average binarized generated image within each of the 11 organelle-defined spatial cluster masks (Clusters 1-11). (3) Fold change (organelle markers): the corresponding spatial enrichment of binarized occupancy within each organelle-defined cluster relative to the mean occupancy across all other non-background clusters, used for the enrichment dot plot in Fig. 5G. (4) Expression (pathway markers): equivalent binarized image intensities computed within the 7 pathway-defined spatial cluster masks (Clusters 1-7). (5) Fold change (pathway markers): the corresponding fold changes for pathway-defined clusters, used for the enrichment dot plot in Fig. S7F.

#### 3 Caption for Video S1.

***ProtiCelli* simulates immunofluorescence stain across hundreds of proteins in a single U2OS cell.** Each frame displays a representative U2OS cell as a 4-channel composite image: the red, blue, and yellow channels correspond to cellular reference markers (microtubules, nucleus, and ER, respectively), and the green channel displays the *ProtiCelli* simulated immunofluorescence stain for a single protein of interest. Proteins are drawn from a curated set spanning major subcellular compartments, including the mitochondria, nucleus, nucleolus, endoplasmic reticulum, Golgi apparatus, cytoskeleton, and plasma membrane. The protein name is shown at the top of each frame, and the frame index is indicated in the lower right corner.
